# Genome-Resolved Metagenomics Analysis of Rice Straw Degradation Experiments Unveils MAGs with High Potential to Decompose Lignocellulosic Residues

**DOI:** 10.1101/2025.03.13.642948

**Authors:** Jeferyd Yepes-García, Nicolás Novoa-Montenegro, Vanessa Otero-Jiménez, Daniel Uribe-Vélez, Emiliano Barreto-Hernández, Laurent Falquet

**Affiliations:** Department of Biology, University of Fribourg, Fribourg, Canton of Fribourg, 1700, Switzerland; Swiss Institute of Bioinformatics, Lausanne, Vaud, 1015, Switzerland; Agricultural Microbiology Group, B Instituto de Biotecnología, Universidad Nacional de Colombia, A.A 14-490, Bogotá D.C., Colombia; Department of Soil and Water Systems, University of Idaho, 875 Perimeter Drive MS2340, Moscow, ID 83844-2340, United States of America; Bioinformatics group, Instituto de Biotecnología, Universidad Nacional de Colombia, A.A 14-490, Bogotá D.C., Colombia

**Author notes:** Correspondence should be addressed to L.F.

## Abstract

**Background:** Rice is one of the top three crops that contribute 60% of the calories consumed by humans worldwide. Nonetheless, extensive rice harvesting yields more than 800 million tons of rice straw (RS) per year globally, generating a byproduct that is often difficult for farmers to manage efficiently without burning it. As a result, millions of tons of carbon dioxide and greenhouse gases are released, causing issues such as respiratory problems, soil degradation, and global warming. In this work, we explore the biological decomposition of RS through the application of microbial consortia from a metagenomics perspective.

**Results:** We applied different treatments to RS placed in a mulching setup during experiments carried out in Colombian rice fields, using various combinations of a *Trichoderma*-based commercial product, the bacterial strain *Bacillus altitudinis* IBUN2717, inorganic nitrogen, and a mixture of potassium-reducing organic acids. Before inoculation and after 30 days of treatment, we characterized the microbial community on the RS surface and from the bulk soil by performing a reference-based compositional analysis, and reconstructing and functionally annotating Metagenome-Assembled Genomes (MAGs). High-quality MAGs with great potential to decompose RS, represented by the extensive number of carbohydrate-active enzymes, were recovered. Soil MAGs taxonomic classification indicates that they may represent potential novel microbial taxa. At the same time, the main part of the RS MAGs with superior lignocellulose-degrading capacity were affiliated under Actinomycetota and Bacteroidota phyla. Moreover, β-glucosidase activity measurements indicated an increased RS degradation after the application of the treatment that included inorganic nitrogen.

**Conclusions:** This contribution underscores the possibility of promoting RS degradation through the application of biological strategies. Further, the newly unveiled MAGs with high RS-degrading potential provide a valuable resource for exploring the functional potential of previously uncharacterized microbial diversity in Colombian agricultural ecosystems, including microorganisms that have not been previously reported as remarkable lignocellulose decomposers.

**Graphical abstract:** 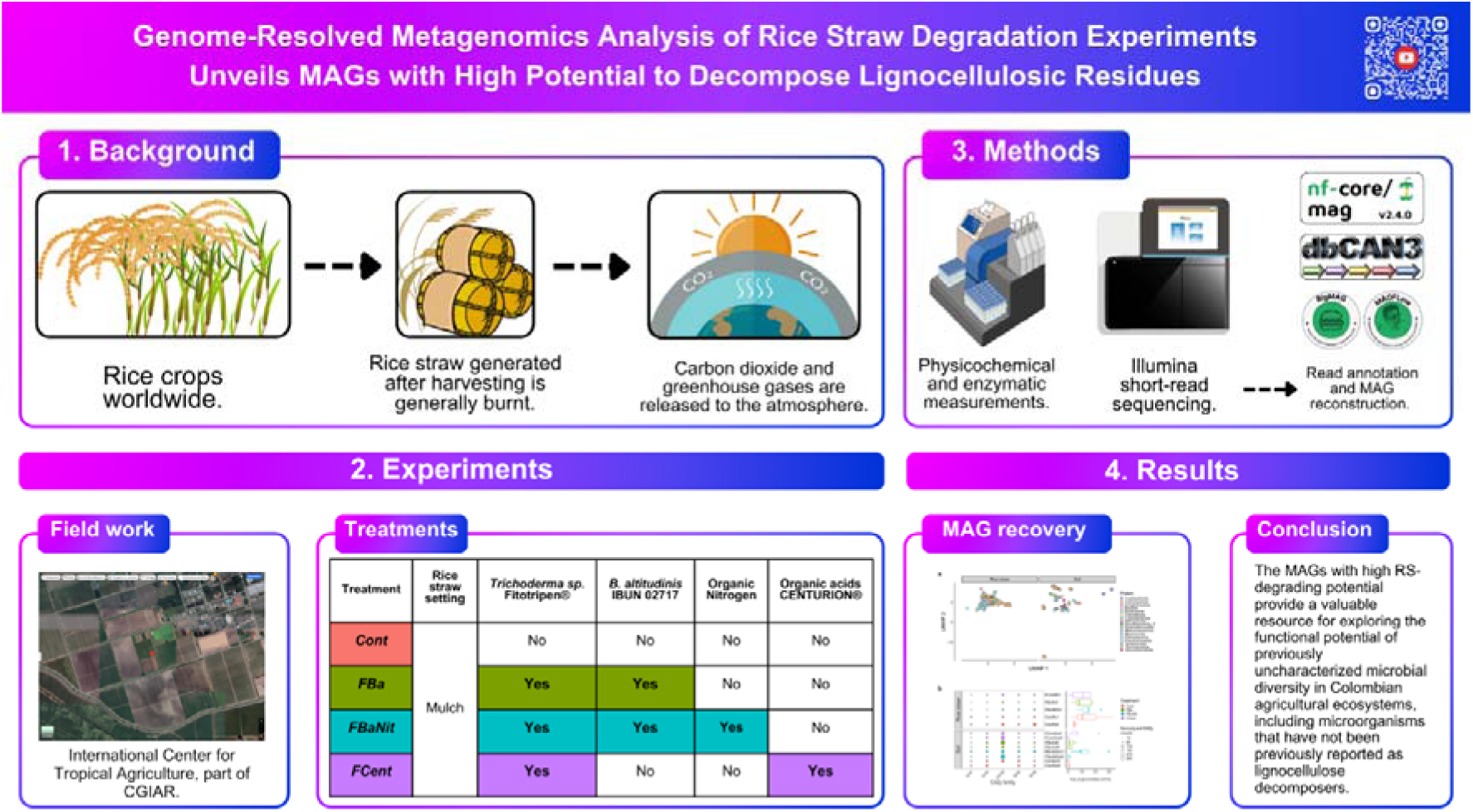

## Introduction

The management of rice straw (RS), a rice agroindustry byproduct, has become one of the most burdensome issues worldwide, challenging researchers to develop innovative and sophisticated strategies to overcome the climate problem RS burning represents [1]. Despite species-specific variations, cellulose (24-38%), hemicellulose (25-27%), lignin (12-13%), plus ashes and silica, make up the constitution of RS, which are organized in a convoluted and inflexible three-dimensional structure [2, 3]; this composition converts RS into a suitable candidate to develop sustainable biologically-based exploitation strategies.

Notwithstanding, the RS degradation, similar to many agro-industrial residues, represents a complex process governed by different factors, including microbial community composition, nutrient availability, and physico-chemical parameters such as pH, temperature, and weather conditions [4]. For instance, the significance of the C/N ratio of organic amendments in controlling process respiration, microbial biomass, and nutrient availability has been described by Kalkhajeh et al. (2021) [5]. Also, the inclusion of blends containing organic acids and organic carbon, i.e., the commercial product Centurion®, during RS bio-transformation contribute to carbon amendment and promote organic matter decomposition [6]. The microbial composition effect on RS degradation has been studied through various strategies, including composting [7], the use of swine-industry waste [8], and the application of commercial products to promote its decomposition [9].

Among several microorganisms documented with the capacity to decompose lignocellulosic matter, *Trichoderma* species stand out as producers of enzymes related to cellulose, hemicellulose, and lignin degradation, as well as biological controllers of common plant pathogens, including the genera *Fusarium*, *Rhizoctonia* and *Botrytis* [10]. Besides enhancing the enzymatic breakdown of RS, *Trichoderma* species can improve the stability of the microbial community during the degradation process [11]. A recent report highlighted how *Trichoderma*-mediated RS decomposition can promote plant growth and impart stress tolerance [12]; whilst, in an interesting study, a setting for solid-state anaerobic digestion of RS that included inoculation with *Trichoderma reesei* improved cellulose degradation after 30 days with a moisture content of 85% [13].

In the case of bacterial strains characterized as lignocellulosic decomposers, the genus *Bacillus* constitutes an interesting option to consider during the execution of experiments aiming at RS degradation, given its high production of a broad range of extracellular enzymes, including cellulases, xylanases, and ligninases [14]. Within *Bacillus*, *B. altitudinis* stands out as a producer of lignocellulose-degrading enzymes, as well as secondary metabolites, including bacteriocins such as pumilarin and altitudin A, contributing to maintaining a balanced and functional microbial community during the decomposition process [15–17].

Examples of treatments involving the blended application of microbial consortia have been published by Cruz-Ramírez et al. (2017) [18], who proposed an improved method for promoting rice-cropping agronomic variables, in which RS was treated with a microbial community composed of *Trichoderma* species, a *Bacillus* strain and small amounts of nitrogen fertilizer. Likewise, Otero-Jiménez et al. (2021) [19] applied a similar strategy, without an inorganic nitrogen fertilizer source, to the RS degradation system at field-scale. In such a study, mulching treatments promoted an abundance of *Acidobacteria* in the rice rhizosphere, whilst non-mulching experiments increased the presence of *Gammaproteobacteria*, *Bacteroidia,* and *Campylobacteria*. Similarly, Kumar et al. (2024) [20] established the high potential of a mixture of fungi (*Aspergillus*, *Trichoderma*, *Fusarium*, and *Rhizopus*) and bacteria (*Pseudomonas* and *Bacillus*) for rapid decomposition of RS through the analysis of Fourier Transform Infrared Spectroscopy (FTIR) spectra. Of particular interest are the cases of Sarma et al. (2022) [21], who enhanced RS composting efficiency up to 87% with the addition of microbial consortia previously isolated within the same work, Qu et al. (2023) [22], whose composting assays conducted to the identification of Actinobacteria, Proteobacteria, Ascomycetes, and Basidiomycetes as driving factors for RS biotransformation, and Ma et al. (2024) [23], who developed straw biodegradation synthetic bacterial consortium following a “top-down” approach that increased cellulose, hemicellulose and lignin conversion up to 66.74_J%, 63.79_J%and 42.85%, respectively.

Furthermore, in order to understand the roles of the members belonging to the microbial community surrounding the decomposition process, data analysis of either 16S rRNA sequencing (amplicon) or whole-genome sequencing (WGS or shotgun) are the common approaches [24]. Although both strategies have their advantages, shotgun sequencing possesses unique features such as a higher taxonomic resolution, a broader coverage of microbial diversity, the detection of novel microorganisms, and the opportunity to recover Metagenome-Assembled Genomes (MAGs), among others [25]. Specifically, by reconstructing the MAGs, it is possible to integrate functional and taxonomic annotation to generate organized and defined models characterizing specific metabolic activity while the process is ongoing [26]. This complementary approach has been successful in recent reports, given the availability of well-organized resources such as the Carbohydrate-Active Enzyme (CAZy) database [27], developed to integrate genomic, structural, and biochemical information on carbohydrate-activated enzymes. For instance, different ecological or experimental environments have been the subject of research during analysis related to the contribution of carbohydrate_Jactive enzymes to marine sediment biodiversity [28], the presence of carbohydrate_Jdegrading enzymes derived from composting samples [29], and the global distribution of carbohydrate utilization potential [30].

Nevertheless, despite the efforts of the aforementioned works to characterize and study the impacts of RS sustainable decomposition on rhizosphere microbiomes and plant growth, the microbial evolution of ongoing RS biotransformation in exterior local conditions remains as an open opportunity to enhance the process. Specifically, taxonomic profiling, metagenome reconstruction, discovery of new native species, and the identification of enzymes involved in lignocellulose breakdown represent pending tasks, especially in Latin American soils.

This study aims to contribute to understanding the microbial community involved in RS degradation in field trials by exploring the application of different treatments encompassing biological, organic and/or inorganic components combined with WGS analysis. As a result, we hypothesized that treatment application would affect the microbial compositionality of the degradation matrices, MAGs with strong lignocellulose-biotransformation abilities can be recovered from RS-degrading experiments at field scale, as well as an enhanced RS degradation considering the reported benefit of the treatment components. To evaluate these hypotheses, we implemented a multi-step bioinformatics pipeline to identify both taxonomic and functional insights from RS-degrading mulching settings, with special interest in MAG reconstruction and characterization of novel microorganisms encompassing a high number and a wide variety of carbohydrate-degrading enzymes.

## Methods

### Area of study

The field assays were carried out in rice fields owned by the Rice Program of the Alliance for Biodiversity and the International Center for Tropical Agriculture (Alliance Biodiversity & CIAT) Km17 straight Cali - Palmira (N3° 29.81539 W76° 21.59739). The plot was divided into and evaluated using furrows measuring 1 m wide by 15.5 m long. The subdivisions were fully randomized to host the different treatments (*see* below), accounting with five replicates per treatment (***Fig. S1***, ***Additional file 1***). Prior to experiment establishment, the rice was harvested by hand and chopped with a scythe.

### Biological material

In order to evaluate the capacity of several biological-based approaches to decompose RS, field experiments were conducted using different combinations of microorganisms and inorganic/organic compounds. The amount of RS used for the experiments was estimated considering an RS accumulation of 3 tons per hectare according to the yield of paddy rice harvested in this location. This estimate was calculated by extrapolating the values given by the collection and weighing of RS contained in a 50 x 50 cm square in five different places in the field. The applied treatments were composed by the addition of some of the following components: *i)* the commercial product Fitotripen^®^ manufactured by Natural Control S.A. (La Ceja, Antioquia Colombia) was utilized; this fungal mixture encompasses three *Trichoderma* strains, namely *T. harzianum*, *T. koningi*, and *T. viridae*, and it is formulated as a powder that must be diluted in water. *ii) Bacillus* strain indexed as IBUN 2717 in the Bacterial Strain Repository of the Biotechnology Institute of the National University of Colombia; this is a native strain isolated from the rice rhizosphere in Colombia, and it has been used in previous studies aiming at biotransform RS [18, 19]. *iii)* Centurion^®^, a carbon amendment for agricultural waste degradation described as an enhancer for RS degradation under field conditions. The *Bacillus* strain required a pre-activation step, where it was grown in Luria Bertani (LB) broth media for 96 hours at 120 rpm at 30°C until its sporulation. The biomass was washed with distilled sterile water (DSW) and heated at 80°C for 10 minutes to eliminate vegetative cells and retain only the spores. The resulting spore suspension was diluted to a concentration of 1×10^7^ spores.mL^-1^ and tested using the MicroDrop technique described by Bautista et al. (2016) [31].

### Field experiments

***Table 1*** describes the composition of the applied treatments as follows: treatment ***FBa***, a suspension was prepared using the *Bacillus altitudinis* strain with Fitotripen^®^ (*Trichoderma* sp.); treatment ***FBaNit***, the same composition as treatment *FBa* in combination with inorganic nitrogen; and treatment ***FCent***, only the *Trichoderma* consortia was used plus 5 mL of the commercial product Centurion^®^ diluted in one liter of distilled sterile water (DSW). The final concentration of the microorganisms was adjusted to 1×10^6^ conidia.mL^-1^ of the *Trichoderma* consortia, and 1×10^7^ spores.mL^-1^ of *B. altitudinis* in a final volume of 1 L in DSW. In the case of treatment *FBaNit*, the inorganic nitrogen was applied in its urea form, aiming to achieve a theoretical carbon/nitrogen (C/N) ratio of 35:1. In consequence, a dose of 56 g of urea (commercial granulate) equivalent to 37.3 kg.ha^1^ of urea was sprayed into the respective plots (15.5 m^2^) after dilution in DSW. Control (***Cont***) plots with the same dimensions as the treatment plots were sprayed with one liter of DSW.

**Table 1.** Treatment composition for the experiments carried out at field scale.

| Treatment | RS setting | Fitotripen® (*Trichoderma* sp.) | *B. altitudinis* IBUN 2717 | Urea | Centurion® (organic carbon and acids) |
| --- | --- | --- | --- | --- | --- |
| Cont | Mulching | No | No | No | No |
| FBa |  | Yes | Yes | No | No |
| FBaNit |  | Yes | Yes | Yes | No |
| Fcent |  | Yes | No | No | Yes |

Following treatment application, the plots were left undisturbed in the field for 30 days under ambient temperature and rainfall. Soil and RS samples were collected at two-time points: before treatment application (time *0*, *t0*) and 30 days later (time *1*, *t1*). At *t0*, the RS samples were taken from a single bulk source, which was then subsampled in specific amounts to initiate the experiments, thus resulting in one sample (with five replicates) at *t0*.

Five spots per plot were selected for soil sampling, each separated by a distance of 2.5 m within each plot (***Fig. S2a***, ***Additional file 1***). Samples were taken using a tube probe (2.54 cm of diameter) that was inserted 5 cm into the soil, and they were placed in a Ziplock® bag and stored in a portable freezer. For RS sampling, two points were selected, each separated by a distance of 5 m from the edge of the plot (***Fig. S2b***, ***Additional file 1***). Before laboratory analysis, soil and RS samples from each replica were combined into a compound sample per plot (biological replica). This was performed in a single run for all samples (time *0* and *1*) to avoid introducing effects related to external variables.

### Soil physicochemical analyses

The field assay comprises physicochemical and enzymatic activity tests. Bulk soil and RS samples were taken from the field as described in ***Fig. S2c-g*** (***Additional file 1***), and frozen at −80°C after arrival to the laboratory. Composite soil samples were prepared, and five replicates per treatment were used to determine pH, total carbon, total nitrogen, phosphate, and enzymatic activity, along with one blank per replicate. The pH was measured in a solution of soil:water in proportion 1:1 according to Miller & Kissel [32], while Dumas dry combustion was used to quantify total carbon and nitrogen in a Total Flash 2000 Elemental Analyzer (Thermo Scientific), and available phosphorus was estimated using the Bray methodology [33]. To test significant differences among treatments, a Shapiro test was implemented to establish the normality of each variable, with a posterior Analysis of Variance (ANOVA) that evaluated the effect of the treatments over individual variables; for visualization purposes, the raw values were transformed using min-max normalization.

### Enzymatic activity

For the enzymatic “cocktail” extraction from the RS samples, 7 g of fresh weighted RS were introduced into a Schott® flask container with 30 ml of sterile saline solution (NaCl 0.85%) and incubated at 30°C temperature for one hour at 200 RPM in an orbital shaker Thermo Fisher^®^ model solaris 4000 [34]. Then, 5 ml of the supernatant was mixed with 100 µl of glycerol and stored at −20°C. After defrosting, protease activity was determined using tyrosine colorimetric estimation with sodium caseinate as a substrate, whilst acid and alkaline phosphatase activities were quantified through the estimation of *p*-nitrophenol (*p*NP) residuals from the interaction between phosphate and phosphatase enzymes during the hydrolysis of phosphate esters of phosphoric acid anhydrides [35]. See ***Additional file 1*** for a detailed explanation in regards to the effect of the treatments on the physicochemical variables, as well as on protease, acid and alkaline phosphatase activity. In addition, β-glucosidase activity was established using a colorimetric procedure that quantifies the residual *p*NP resulting from the enzyme’s interaction during the hydrolysis of glycosides [35]. Similar to the soil physicochemical analysis, through a Shapiro test we determined the normality of the data, and significant differences among treatments were screened by running a Kruskal-Wallis test with a consequent Dunn test if required.

### DNA extraction and sequencing

RS DNA isolation was conducted in accordance with the methodology delineated by Su et al. (2022) [36]. The procedure entailed washing 1 gram of straw in a PBS buffer at 120 rpm for one hour, followed by two sonication sessions for one minute at 30 kHz. The resulting liquid PBS suspension was subsequently subjected to centrifugation at 15,000 rpm for 10 minutes, resulting in the isolation of an RS pellet. For soil and RS pellet Power Soil Pro Kit® was employed. The concentrations and quality of the extracted DNA were then estimated using a Qubit and gel electrophoresis. A mock community (ZymoBIOMICS™ Microbial Community DNA Standard D6305) was included as an additional sample during the extraction and sequencing procedures to check the quality of both experimental and computational workflows.

### Reference-based compositional analysis of Next-generation sequencing data

The quality of the raw sequences was established using FastQC v0.12.1, and the reports were concatenated by MultiQC v1.16 [37]. After sequence integrity verification, genome host removal was carried out by sequence alignment with the indexed genome of rice by Bowtie2 [38] (*O. zativa*, NCBI RefSeq accession number: GCF_000005425.2); the unaligned reads were taxonomically annotated with Kraken2 [39] v2.1.3 relying on database PlusPFP (Standard plus RefSeq protozoa, fungi & plant, released on 04.09.2024). A Bayesian species abundance re-estimation was performed with Bracken [40] v2.8 to finally generate interactive metagenomic visualizations through Krona plots [41]. This described workflow was proposed by Lu et al. (2022) [42], and it was wrapped as a Nextflow (DSL2) [43] pipeline to enhance the reproducibility and portability of the analysis.

The resulting Bracken files were merged in BIOM format [44] to generate a Phyloseq [45] v1.42.0 object afterwards. Relative abundance was estimated at domain level for merged, agglomerated at this taxonomic level, samples of RS and soil matrices. Whilst at phylum and genus relative abundance was measured for merged replicates of each sample; at each level, the data were agglomerated correspondingly.

Following the abundance analysis, the counts were transformed into Counts Per Million (CPM) as a normalization method to mitigate the effect of the size library on the analysis, and given that the domain *Bacteria* represented more than 95% of the annotated reads, the diversity analysis was performed only with the reads belonging to this domain. α-Diversity was measured as Shannon and Inverse Simpson indices, while β-diversity was assessed with Principal Coordinate analysis (PCoA) on Bray-Curtis distances and through Uniform Manifold Approximation and Projection (UMAP) on Aitchison distance (Euclidean distance on centered log-ratio transformed data) in order to count with both parametric and non-parametric estimations. In addition, compositional variability was tested by measuring the distance to the centroid of the multivariate space obtained with the Aitchison distance, along with an ANOVA to test the significance of the distances. Furthermore, significant differences in microbial composition were tested using Permutational Multivariate Analysis of Variance (PERMANOVA) considering the proportion of the variance explained by the factors Time and Treatment; several dissimilarity/distance metrics (Aitchison, Bray-Curtis, Euclidean and Jaccard) were used as input to verify the consistency of the analysis.

Since RS and soil compositions were significantly distant, the differential abundance analysis with ANCOM-BC2 [46] v2.0.3 was carried out separately for each matrix. The *Cont* samples taken at time *0* were used as a pre-specified group for each case, *p*-values were adjusted via Holm–Bonferroni method, the fixed formula considered Treatment plus Time in the case of soil samples, while only Time or Treatment were set in the case of RS samples; the remaining parameters of the ANCOM-BC2 function were left in their default state. The input data was agglomerated at genus level and filtered for genera present in at least 0.5% in any sample. The downstream analysis was performed in an R v4.2.3 environment that included the packages Vegan [47] v2.6.4 and UMAP v0.2.10.0. The corresponding results from this reference-based compositional analysis, along with a comprehensive discussion about the effect of the treatments on the taxonomic profiles, can be found on the ***Additional file 1***.

### Metagenomic assembly and MAG reconstruction

Initial bins were built using the nf-core/mag [48] v2.3.2 from raw sequences. The following parameters were included during nf-core/mag launching: `--busco_auto_lineage_prok --skip_spades -- host_genome IRGSP-1.0 --skip_concoct --megahit_options ‘--min-contig-len 1000’`. A co-assembly/co-binning strategy was chosen as an attempt to improve the quality and resolution of the bins [49], and hence all replicates belonging to the same sample were processed together. MEGAHIT [50] v.1.2.9 was used for the assembly, while MaxBin2 [51] v2.2.7 and MetaBAT2 [52] v2.15 were the selected tools for binning. A de-replication and refinement step was carried out by enabling DASTool [53] v1.16 within the workflow (`--refine_bins_dastool`). The depth data of each bin provided by the nf-core/mag were considered to estimate their relative abundance and as complement to bin-quality analysis.

Once the bins were generated, a downstream study was performed leveraging on the quality metrics generated by the pipeline MAGFlow/BIgMAG v1.1.0 [54]; this workflow includes quality-measuring tools such as CheckM2 [55] v1.0.1, BUSCO [56] v5.7.0, GUNC [57] v1.0.6 and QUAST [58] v5.2.0, as well as the taxonomic annotator GTDB-Tk2 [59] v2.4.0. The versions of the databases required by this pipeline were Genome Taxonomy Database (GTDB) r220, GUNC reference database (based on proGenomes 2.1) and CheckM2 uniref100.KO.1. Mid and high-quality MAGs were filtered for subsequent analysis using the following criteria [60]: mid-quality MAGs (MQ) with completeness >= 50% and contamination < 10%, and completeness > 90% and contamination < 5% for high-quality (HQ) genomes. Moreover, MAGs belonging to either RS or soil samples with Average Nucleotide Identity (ANI) < 5% were clustered using FastANI [61] v1.33; the representative MAG from each cluster was selected based on the score formula proposed by Rühlemann et al. (2022) [62]: completeness – 0.5 x contamination.

The resulting MAGs were annotated by Prokka [63] v1.14.6, and the coding sequences from each MAG were then subjected for Cluster of Orthologous Genes (COGs) detection by COGclassifier v1.0.5 and Marker genes (MG) annotation with FetchMG [64] v1.3. Similarly, the presence of carbohydrate-related metabolism enzymes (CAZyme modules) was studied through dbCAN3 [65] v4.1.4; a module was considered as present if it was reported by either Diamond alignment or HMMER biosequence analysis. CAZyme modules related to lignocellulosic biomass decomposition (including β-glucosidase enzymes) were established as followed by Santos-Pereira et al. (2023) [29], and additional CAZyme families such as cellobiose-dehydrogenases, laccases and lignin-peroxidases were included. A normalization procedure of the CAZyme counts per MAG across samples was carried out as implemented by López-Sánchez et al. (2024) [28] with the equation:

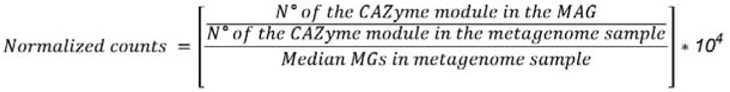

Additionally, to normalize the CAZyme counts per sample, without considering the specific abundance in each MAG, the following formula was used:

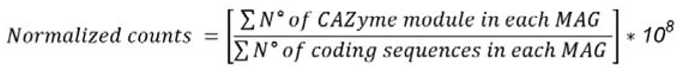

Finally, a concatenated taxonomic tree of the MAGs belonging to each matrix was built using GTDB- Tk2 through the command ‘`gtdbtk de_novo_wf’` and the previously generated taxonomy classification via MAGFlow. Further, this information was cross-referenced with quality metrics data, COG annotation and CAZyme normalized counts to integrate the activity analysis of each genome. The CAZyme normalized-count table was used as input for the UMAP methodology as an attempt to identify clusters of MAGs sharing carbohydrate-related overall presence of enzymes. ***Fig. 1*** introduces the bioinformatics workflows followed above-described in this work to carry out the taxonomic profiling of the raw sequences, and to reconstruct and annotate the MAGs. For visualization purposes, a variety of plots were generated using Circos Table Viewer [66] v0.63-10, iTOL [67] v7.0 and R in-house scripts that include ggplot2 v.3.5.1

**Fig. 1.**
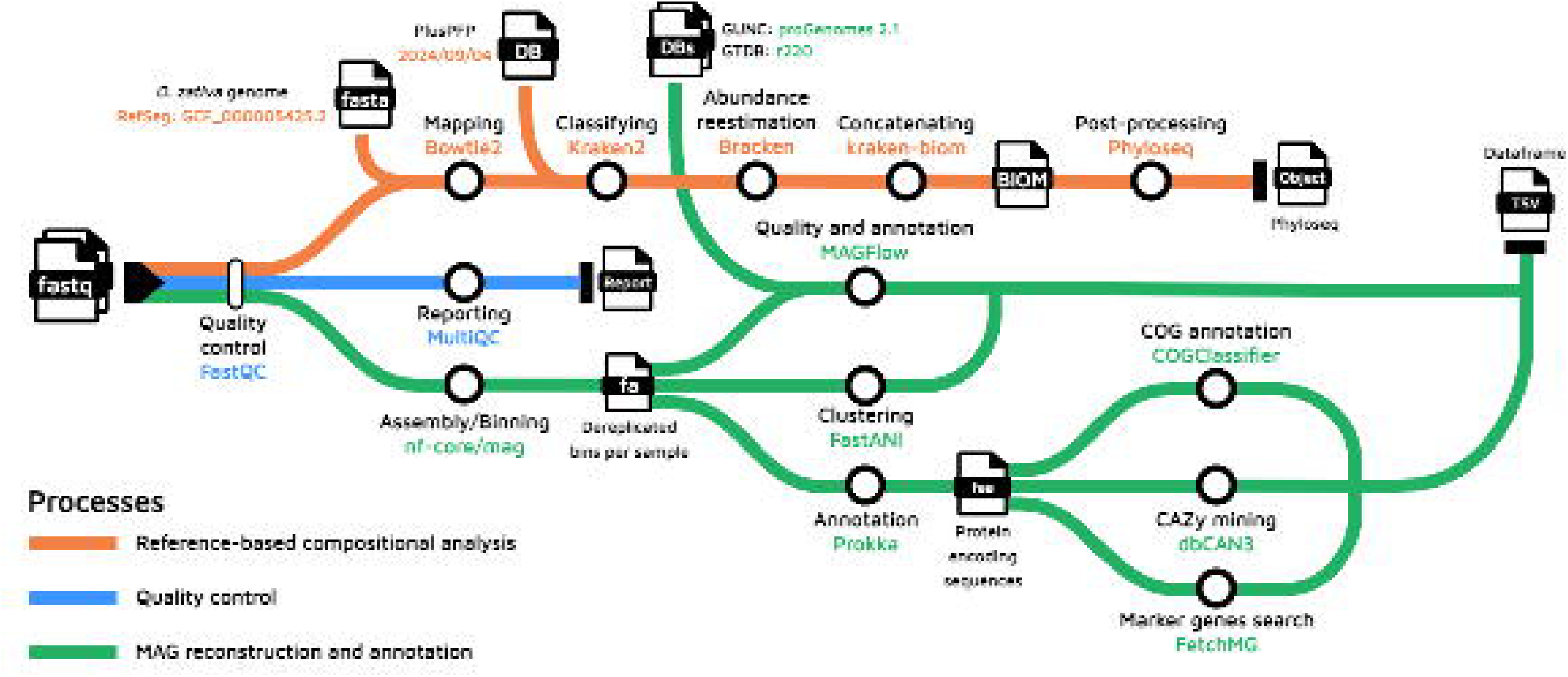
Bioinformatics workflow followed to evaluate sequence quality, perform reference-based compositional analysis, and MAG recovery.

## Results

### MAG reconstruction and annotation

Initially, the bins recovered by nf-core/mag using the co-assembly/co-binning approach were filtered using the MIMAG criteria afore-mentioned, yielding between 28 and 50 MAGs per sample in RS, and between 12 and 19 in soil samples (***Fig. S10***, ***Additional file 1***). In terms of quality, the percentage of HQ MAGs was variable (between 20% and 45%) across samples and across matrices, with a similar trend in the case of the percentage of MAGs passing the GUNC test (> 60% in RS MAGs and > 75% in soil MAGs). In addition, the percentages of deepest annotated taxon were similar for all samples in RS, as well as for all samples belonging to the soil matrices. Notwithstanding, when reviewing the percentage of deepest annotated taxon, soil samples stand out as for none of the MAGs the species level was identified, and even in some cases the annotation was only possible until order level. Remarkably, the fact that GTDB contains more than 580,000 genomes (r220) highlights the intrinsic potential of environmental niches such as the bulk soil or RS considered in this study to contain novel species, and to act as a reservoir for genes with a wide variety of metabolic activity. This situation added to the remaining knowledge gap for more species in the database, accounts for the inability by GTDB-Tk2 to properly classify several MAGs. Likewise, analogous to the read-based compositional analysis, the complexity of the soil community difficulties the assembly process, resulting in a considerably lower number of MAGs built by the pipeline. An additional result description in regards to presence/absence and abundance of the MAGs per sample can be found on the ***Additional file 1***, and complete taxonomic annotations of each MAG belonging to RS or soil samples are available on ***Table S3*** and ***Table S4*** (***Additional file 2***), respectively.

### MAGs and integration of carbohydrate-active enzyme information

In order to identify microorganisms with the highest likelihood of acting as lignocellulosic decomposers, the recovered MAGs were analyzed to detect carbohydrate-related enzymes (CAZymes) and Cluster of Orthologous Genes (COGs). In regard to COGs, matrix-wise (***Fig. S12***, ***Additional file 1***), despite some specific MAG-related variations, the classification distribution of COGs per sample is similar among them to a wide extent; this scenario is persistent across RS and soil samples. Further, the COG category with the highest proportion within the sample coding sequences is Metabolism, followed by Information Storage and Processing, and Cellular Processes and Signaling, implying a potential engagement in processing organic matter.

Moreover, the dimensionality of the normalized CAZyme counts matrix (before MAG ANI-based clustering) was reduced through UMAP graph representation as an attempt to identify groups of MAGs closely related in terms of their potential to decompose cellulose and lignin (***Fig. 2a***). At first glance, although this clustering technique does not separate the MAGs in function of their sample precedence, it truly generates sets of MAGs with similar CAZyme patterns and inclusion within the same phylum mainly in soil samples. This is especially noticeable in the case of soil MAGs, where MAGs belonging to the phyla Acidobacteriota and Actinomycetota aggregate in their respective clusters. In the case of RS samples, the clusters overlapped each other at a central clump, albeit it is still possible to recognize some independent groups of MAGs mostly for Pseudomonadota and Actinomycetota phyla.

**Fig. 2.**
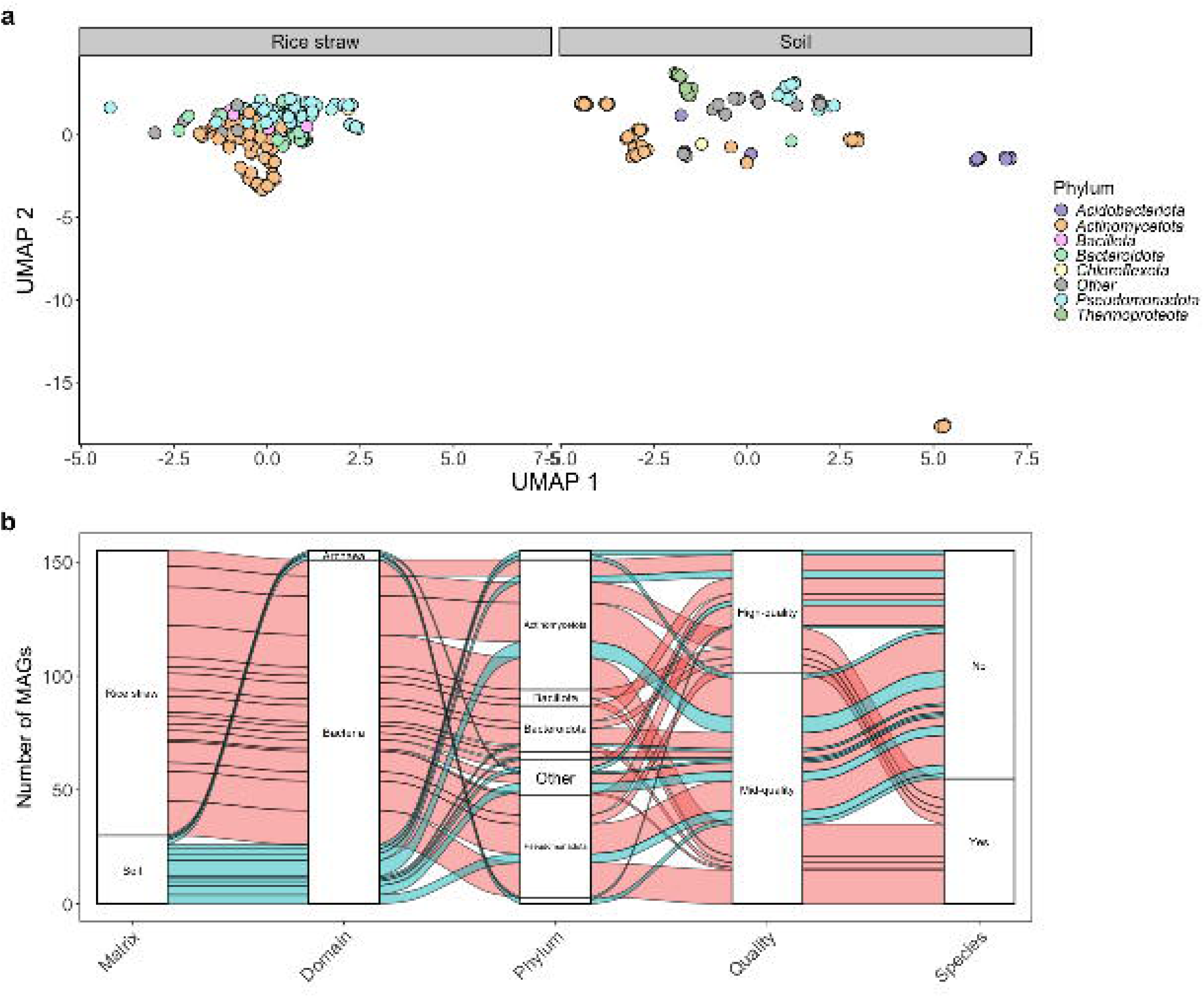
***a*** Two-dimensional Uniform Manifold Approximation and Projection (UMAP) embedding of normalized CAZy enzymes counts per MAG (before clustering with FastANI) colored by phylum (GTDB-Tk2). All CAZy classes (GH, GT, PL, CE, AA and CBMs), along with all their families, were considered for clustering using default values (15 neighbors and Euclidean distance). ***b*** Taxonomic classification (at the domain and phylum levels), quality (CheckM2), and identification as species or not of the clustered MAGs from the RS and soil metagenomes. The exact order of the missing phylum labels within the ***b*** plot can be read from the legend on ***a***. Phylum *Other* includes: Armatimonadota, Cyanobacteriota, Deinococcota, Desulfobacterota_E, Gemmatimonadota, Methanobacteriota, Myxococcota, Patescibacteria, Tectomicrobia and Verrucomicrobiota.

Afterwards, given that substantial differences among treatments in terms of CAZyme general patterns were not detected (***Fig. S11***, ***Additional file 1***), we shifted to a matrix-based analysis by clustering all MAGs belonging to either RS or soil matrices using ANI, and a representative member of each cluster was selected based on their quality score (completeness – 0.5 x contamination); condensed information per representative MAG about taxonomic annotation and functional features is presented on ***Fig. 2b*, *Fig. 3*** and ***Fig. 4***. A total of 125 MAGs were recovered in RS samples, with phyla Pseudomonadota, Actinomycetota and Bacteroidota composing more than 80% of the taxonomic annotation (***Fig. 2b***); the proportion of HQ MAGs was around 37%, and about 80% of the RS MAGs passed the GUNC test. Also, genome size presents a high variability, covering from 1 Mbp to 10 Mbp, even among MAGs belonging to the same phylogenomic cluster (***Fig. 3***). Also, it is important to mention that though it seems that a higher number of MAGs were recovered at time 1, this is a consequence of the experiment setup, in which at time *0* only 5 replicates of the original RS bulk were sequenced, and the MAG sample precedence is due to the clustering and scoring step, and some MAGs can be found in more than one treatment (*see **Fig. S13***, ***Additional file 1***).

**Fig. 3.**
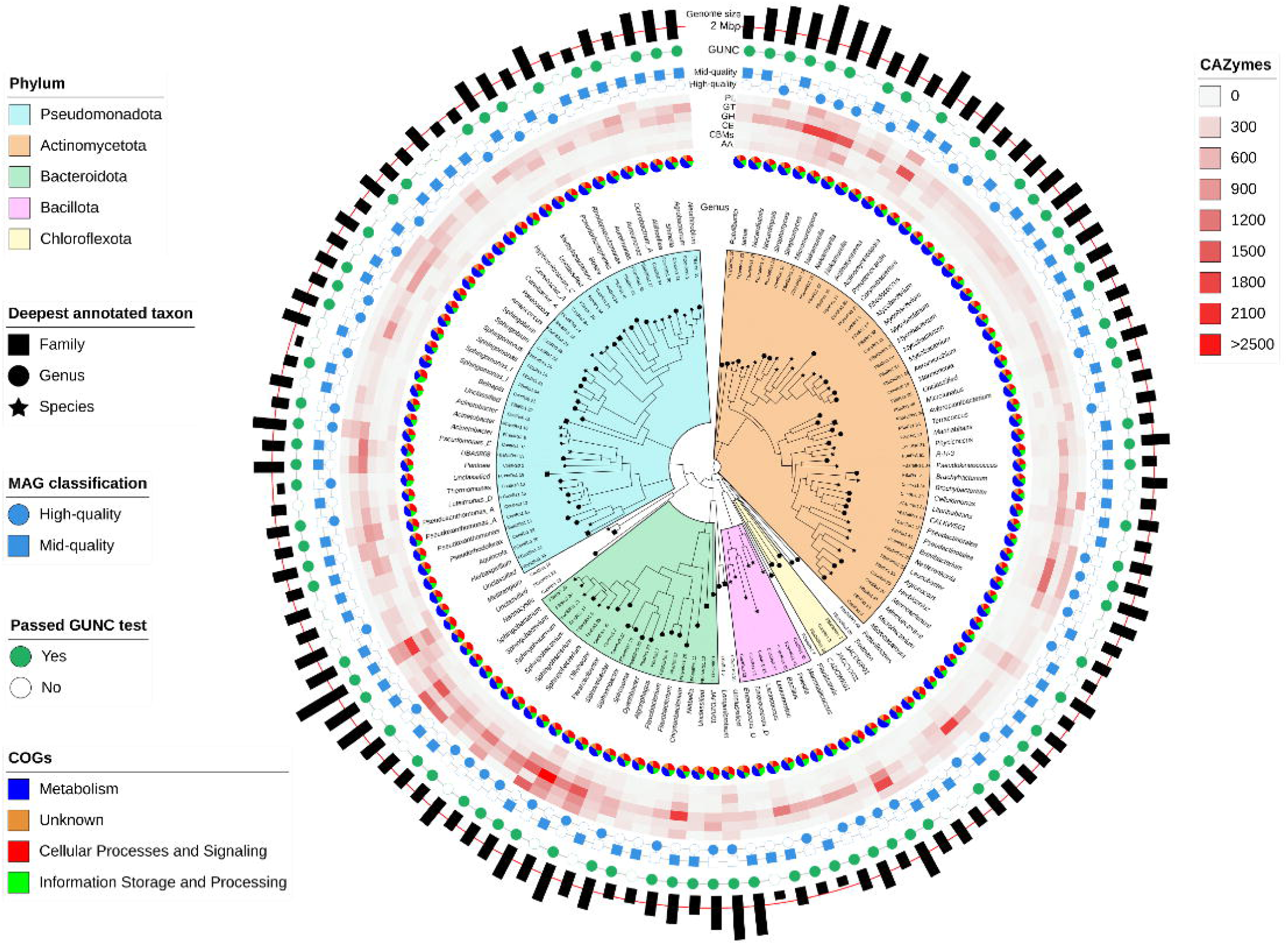
125 recovered MAGs from **RS** samples after clustering based on Average Nucleotide Identity (ANI). The visualization includes phylogenetic relationship among them (tree), specific highlighted phylum clades (colored sectors), genus classification (outer text), proportion of the main categories of Cluster of Orthologous Genes (COG) within each MAG (blue-orange-red-green pie chart), Carbon Active EnZyme (CAZyme) class normalized abundance (red scaled heatmap), classification as either high-quality or mid-quality MAG based on checkM2 measurements (blue binary), GUNC test passing (green binary), and genome size in Mbp (outer black bars).The displayed MAGs are representatives from the generated clusters using ANI, and they were selected based on the score: completeness – 0.5 x contamination. Conventions in CAZyme heatmap: GH glycoside hydrolases, GT glyco-syltransferases, PL polysaccharide lyases, CE carbohydrate esterases, CBMs carbohydrate-binding modules, AA auxiliary activities.

**Fig. 4.**
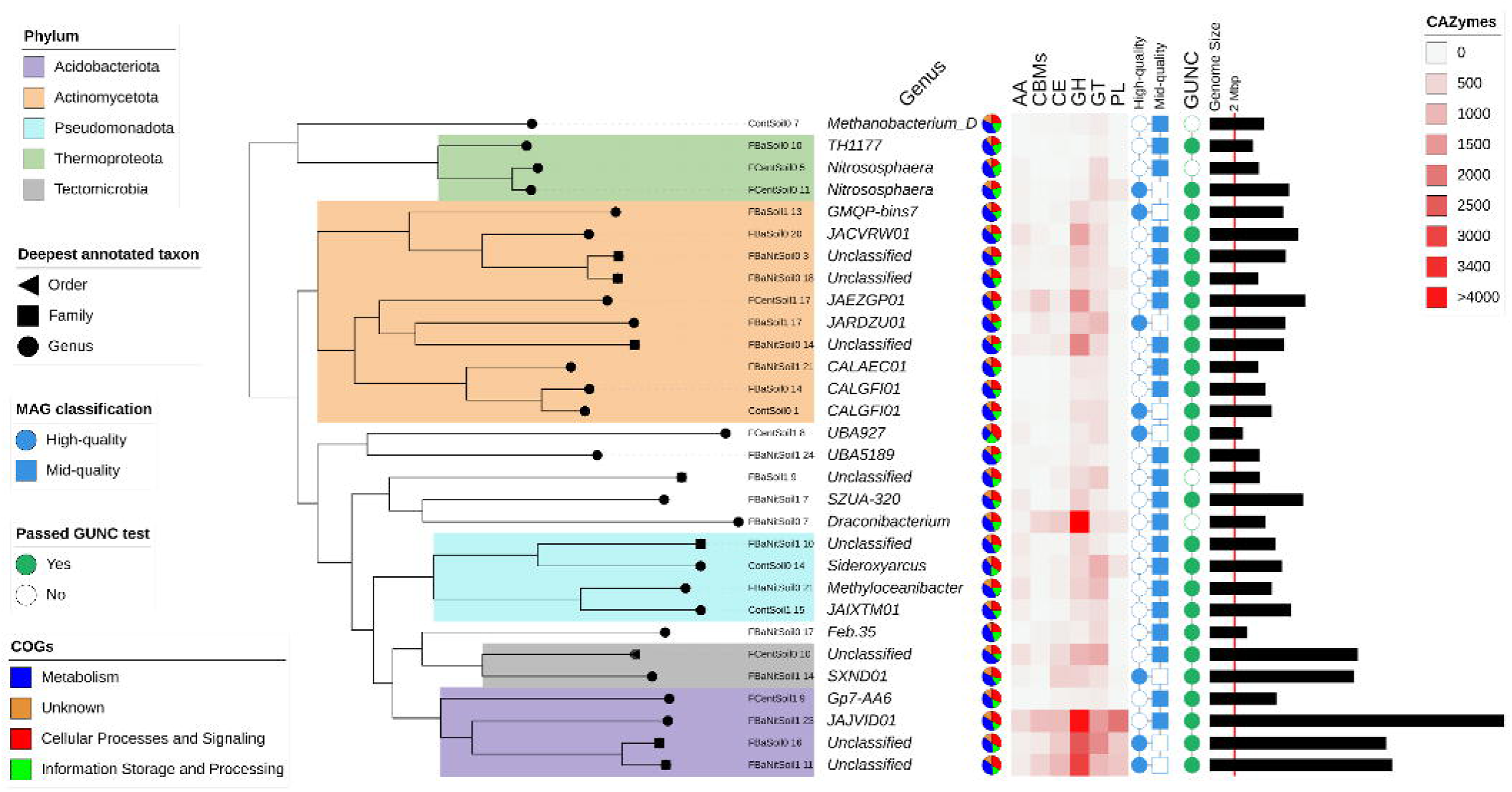
30 recovered MAGs from **soil** samples after clustering based on Average Nucleotide Identity (ANI). *See* ***Fig. 3*** for feature explanation.

Furthermore, in terms of the CAZyme distribution across MAGs, polysaccharide lyases (PLs), glycosyltransferases (GTs) and glycoside hydrolases (GHs) are concentrated mainly on members of the Bacteroidota phylum, specifically in MAGs annotated as *Sphingobacterium*, *Siphonobacter*, *Spirosoma* and *Pararcticibacter*. *Mellitangium* genus constitutes an interesting case as it holds a high prevalence of the previously mentioned CAZyme categories plus carbohydrate esterases (CEs), carbohydrate-binding modules (CBMs) and auxiliary activities (AAs). Remarkably, some Actinomycetota genera such as *Streptomyces* and *Micromonospora* depict a high concentration of the GH modules.

On the other hand, soil samples offer a completely different overview since solely 30 MAGs are found after clustering (***Fig. 4***). Besides some features already described during the MAG presence/absence analysis (*see **Fig. S14***, ***Additional file 1***), the clustered MAGs display a low percentage of HQ MAGs (around 26%), albeit a high percentage of MAGs passing the GUNC test (85%) was observed. The genome size feature highlights MAGs ranging from 2 Mbp to 12 Mbp, where the MAG annotated as the genus *JAJVID01* (p Acidobacteriota) is located at the upper limit in terms of regular bacterial genome size. Strangely, although this MAG passes the GUNC test and it is classified as a MQ MAG, it has twice the genome size of the reference genome in the GTDB, creating some interesting questions in regards to the scope of the quality metrics provided by the software commonly used to establish such criteria. Nevertheless, the phylum Acidobacteriota concentrates the highest presence of CAZymes in all categories, namely GHs, GTs, PLs, CEs, CBMs and AAs, composing probably the phylum in which the carbohydrate-degrading activity relies on. This high abundance of CAZymes within this phylum is interestingly correlated with the highest genome sizes found in the soil MAGs, and it is worth mentioning that MAGs belonging to Acidobacteriota were the most abundant MAGs with at least 30% of the total abundance (see ***Fig. S14***, ***Additional file 1***). Special mention should be addressed to the MAG classified as *Draconibacterium* given that, along with *JAJVID01*, it depicts a high concentration of GHs, the module in which the majority of enzymes related to lignocellulosic metabolism are encompassed.

An immersion into the annotation of GHs families in each representative MAGs is enabled by reviewing the bubble plot depicting the CAZyme families associated with lignocellulose-degrading enzymes families found in the MAGs recovered from RS and soil samples (***Fig. 5*** and ***Fig. 6***); a complete overview of the selected CAZymes per RS MAG can be found on ***Fig. S15*** (***Additional file 1***).

**Fig. 5.**
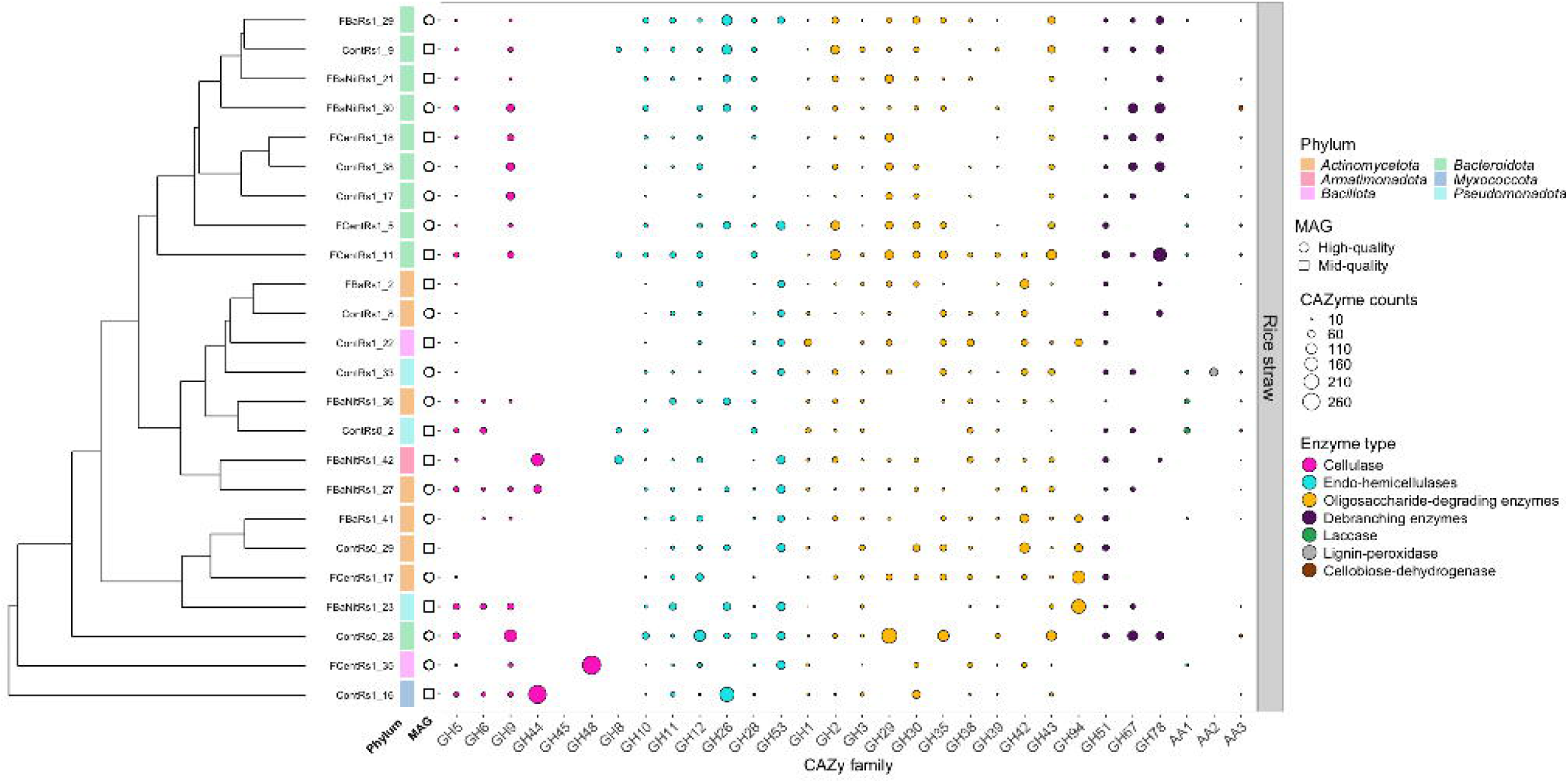
Bubble plot depicting the number of normalized counts for CAZyme families associated with lignocellulose-degrading enzymes found in the MAGs recovered from **RS** samples. Visualization includes: MAG clustering based on normalized CAZyme counts of the selected families, and phylum information (GTDB-Tk2). The classification as MQ MAG or HQ MAG (CheckM2) is designated by the shape on the left. The displayed MAGs are representatives from the generated clusters using ANI, and they were selected based on the score: completeness – 0.5 x contamination. Displayed RS MAGs contain at least 30 normalized counts in 5 or more CAZyme families; complete information about normalized CAZyme counts in RS MAGs is found in ***Fig. S15*** (***Additional file 1***).

**Fig. 6.**
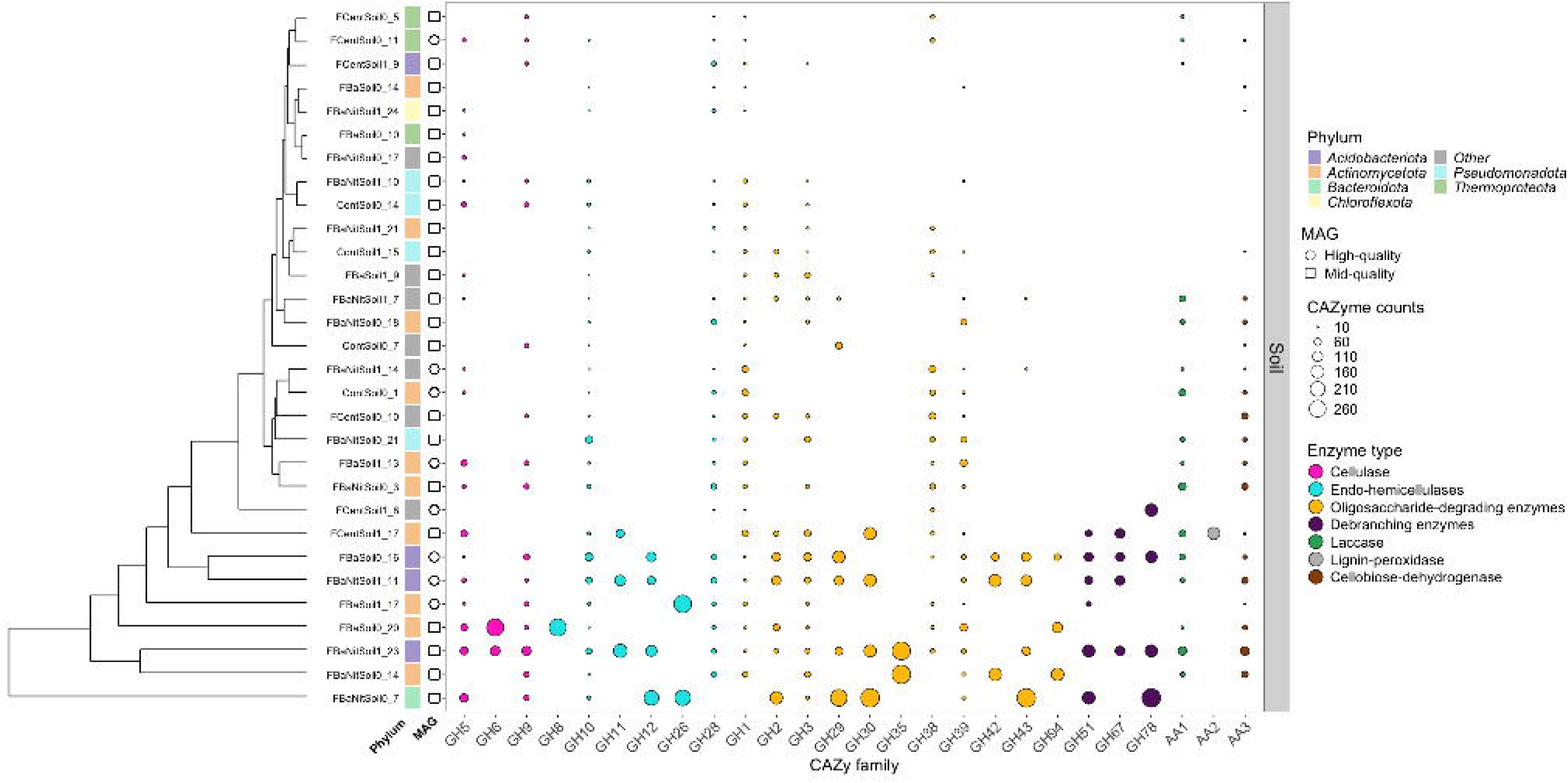
Bubble plot depicting the number of normalized counts for CAZyme families associated with lignocellulose-degrading enzymes found in the clustered MAGs recovered from **soil** samples. *See* ***Fig. 5*** for feature explanation. All soil MAGs are displayed.

Notably, it is important to mention the co-presence within single MAGs of the majority of the selected lignocellulolytic enzymes (considering all of the categories), assigning them a theoretical capacity to decompose agricultural subproducts, including RS, almost in an isolated manner (***Fig. 5***). This scenario would suggest the presence of an established functional redundancy within the RS community regarding metabolizing specific substrates, aiding in the maintenance of functional diversity. Representative examples of these “totipotential” MAGs able to carry out the main part of lignocellulose degrading are *FCentRs1_11* (*Niabella*) and *FBaNitRs1_27* (*Micromonospora*). However, some MAGs seem to be more specialized in specific tasks within the decomposing process, including *FCentRs1_35* (*Bacillus*) and *ContRs1_16* (*Melittangium*) which are presumably more specialized in cellulose rupture given its high abundance of the CAZyme family GH44 and GH48, respectively. A Noteworthy particular case is the MAG *ContRs1_19* (*Nannocystis*), a carrier of multiple copies of genes coding for GH45 (cellulose endo-β-1,4-glucosidase/xyloglucan endo-β-1,4-glucosidase) (***Fig. S15***, ***Additional file 1***).

Complementary, the afore-mentioned MAG *ContRs1_16*, as well as *ContRs0_28* (*Siphonobacter*), *FBaRs1_29* (*Flavobacterium*), and some more, account with a broad range of endo-hemicellulases involved in the initial hemicellulose cleavage. These microorganisms could be complemented by *ContRs1_38* (*Sphingobacterium*), *FCentRs1_18* (*Siphonobacter*) and *FBaNitRs1_30* (*Spirosoma*) based on their high theoretical presence of debranching enzymes (GH51, GH67 and GH78).

Meanwhile, *ContRs0_28*, *FCentRs1_17* (*Pseudodactinotalea*), *FCentRs1_11* (*Niabella*), among others, appear to be more adapted to perform the cellulose/hemicellulose downstream metabolism process considering their variety and abundance of oligosaccharide-degrading enzymes (GH29, GH35, GH38, GH94, among others). On the contrary, the lignin degradation is apparently more scarcely performed as laccases, AA1 family are present in a lower percentage of the MAGs, from which the already mentioned *FBaRs1_2* and *FBaNitRs1_21*, plus *ContRs0_2* (*Pantoea*), *FBaNitRs1_36* (*Streptomyces*) and *ContRs1_33* (*Sphingobium*) stand out. Likewise, lignin-peroxidase families AA2 are carried mainly by *ContRs1_33*, *FBaNitRs1_39* (Unclassified genus) and *FCentRs1_23* (*Iamia*).

Correspondingly, among the soil MAGs (***Fig. 6***), it is also possible to identify MAGs able to metabolize cellulose, hemicellulose and lignin almost in isolation such as *FBaNitSoil1_11* (p Acidobacteriota, genus unclassified), *FBaNitSoil1_23* (p Acidobacteriota, g_*JAJVID01*) and *FBaSoil0_16* (p Acidobacteriota, genus unclassified), suggesting a kind of specialization on this particular substrate within the microbial community. It is worth mentioning how the MQ MAG *FBaNitSoil0_7* (*Draconibacterium*) surprisingly appeared to contain a high number, and a wide variety, of genes that would be potentially expressed as oligosaccharide-degrading enzymes, endo-hemicellulases, debranching enzymes and cellulases. Also, *FBaSoil0_20* (p Actinomycetota; g *JACVRW01*) and *FBaNitSoil1_23* exhibit an important proportion of endo-β-1,4-glucosidases (GH5, GH6, GH9), being probably the members of the community associated with the bioconversion of cellobiose into glucose. Similarly, endo-hemicellulases are present in several MAGs, although a small set of MAGs seemed to concentrate the greatest numbers of the CAZymes, namely *FBaNitSoil1_23*, *FBaSoil0_20, FBaNitSoil1_11*, *FBaSoil0_16* and *FCentSoil1_17* (p Actinomycetota, g *JAEZGP01*). Equally, oligosaccharide-degrading enzymes were clustered mostly within *FBaSoil0_16, FBaNitSoil1_11, FBaNitSoil1_23, FBaNitsoil0_14* (p Actinomycetota, genus unclassified) and *FBaNitSoil0_7*; these same MAGs, excepting *FBaNitsoil0_14,* retain the main part of the debranching enzymes. Nonetheless, in contradistinction to RS MAGs, lignin-degrading CAZymes (AA1) are widespread across more than a half of the MAGs, while AA2 family (lignin-peroxidase) appeared to be present solely in a small number of MAGs, being *FCentSoil1_17* the principal carrier of this type of enzymes.

Likewise, and as mentioned previously for RS MAGs, within soil MAGs, there seems to be microorganisms particularly in charge of certain decomposition tasks, which suggests as well the configuration of functional redundancy within the microbial community. In this case, it is noticeable how elevated counts of GH35 (oligosaccharide-degrading), GH6 (cellulases), GH8 and GH26 (endo-hemicellulases) were detected only in one or two soil MAGs. An analogous case was detected in regards to GH45 and GH48 (cellulases) were present in only one RS MAG.

After grouping the CAZyme annotation of RS MAGs and soil MAGs independently (***Fig. 7***), it is noticeable how certain enzyme families are overrepresented within each matrix. For instance, AA3 (module including cellobiose dehydrogenase), GH1 (cellulases), and GH10 (endo-hemicellulases) were observed as the most abundant families within the MAGs belonging to both matrices; whereas GH43 and GH3 (oligosaccharide-degrading) were remarkably abundant in RS samples. This high number of specialized enzymes may indicate that the biotransformation of RS was active regarding two of the main components of this residue, cellulose and hemicellulose.

**Fig. 7.**
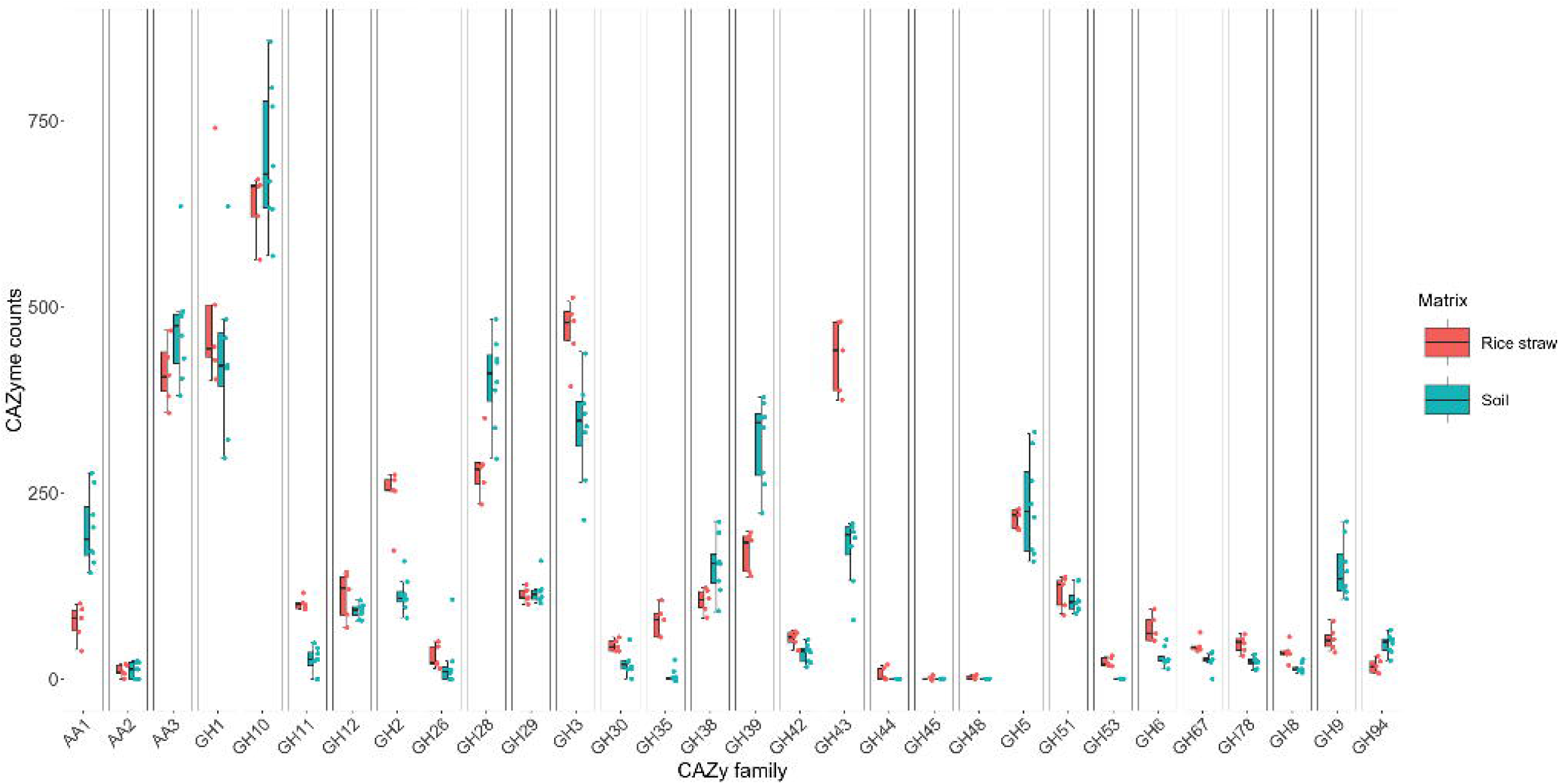
Boxplot showing the distribution and abundance of normalized counts for CAZyme families associated with lignocellulose-degrading enzymes found in the MAGs recovered from RS and soil samples. Values displayed are the cumulative sum of each CAZyme family within a determined sample, normalized by the number of coding sequences found in the sample.

The analysis of the normalized CAZyme counts (***Fig. 7***) showed that certain enzyme families were more abundant in one matrix compared to the other. Of particular interest is the number of enzyme families that are relatively more abundant in the soil matrix than in the RS (AA1, GH28, GH39, and GH9), suggesting that microbial taxa harboring these enzymes are better represented in the soil matrix. This may indicate that the application of RS treatments could stimulate their expression. Similarly, regardless of the RS treatment, the enzyme families GH2, GH3, GH11 and GH43 are more prevalent in the RS matrix, highlighting that a higher number of taxa encompassing these enzyme families exist in the RS than in the soil. These results highlight differences in enzyme family abundance, likely due to specific niche specializations. Further studies could analyze these findings in greater depth to assess their functional implications.

### βlZGlucosidase experimental activity in RS and soil samples

Given that GHs have a significant capacity for the breakdown of plant biomass, specifically cellulose and hemicellulose, and are one of the primary CAZyme families found in RS and soil samples, we performed an experimental validation of the presence of specific members of these enzymes within the samples. Specifically, we measured the βIi.Glucosidase activity as this enzyme is usually a limiting factor during cellulase biotransformation and glucose releasing. We observed an increase in the β-glucosidase activity in both matrices, which indicates an increment in the cellulose biotransformation after 30 days of the treatment application (***Fig. 8***). RS samples showed up to 5-fold of augmented enzymatic activity once finished the experiments, including the *Cont* treatment. An analogous, and more pronounced, scenario is depicted by soil samples, where treatments *FBaNit* and *FCent* present activity values up to 20 times higher after 1 month compared to the initial measurements. Notwithstanding, significant differences among treatments were only detected in soil samples, where the treatment *FBaNit* demonstrated a significant increase of the βIi.Glucosidase compared to *Cont* samples after 30 days of assay (*see **Table S5***, ***Additional file 2***).

**Fig. 8.**
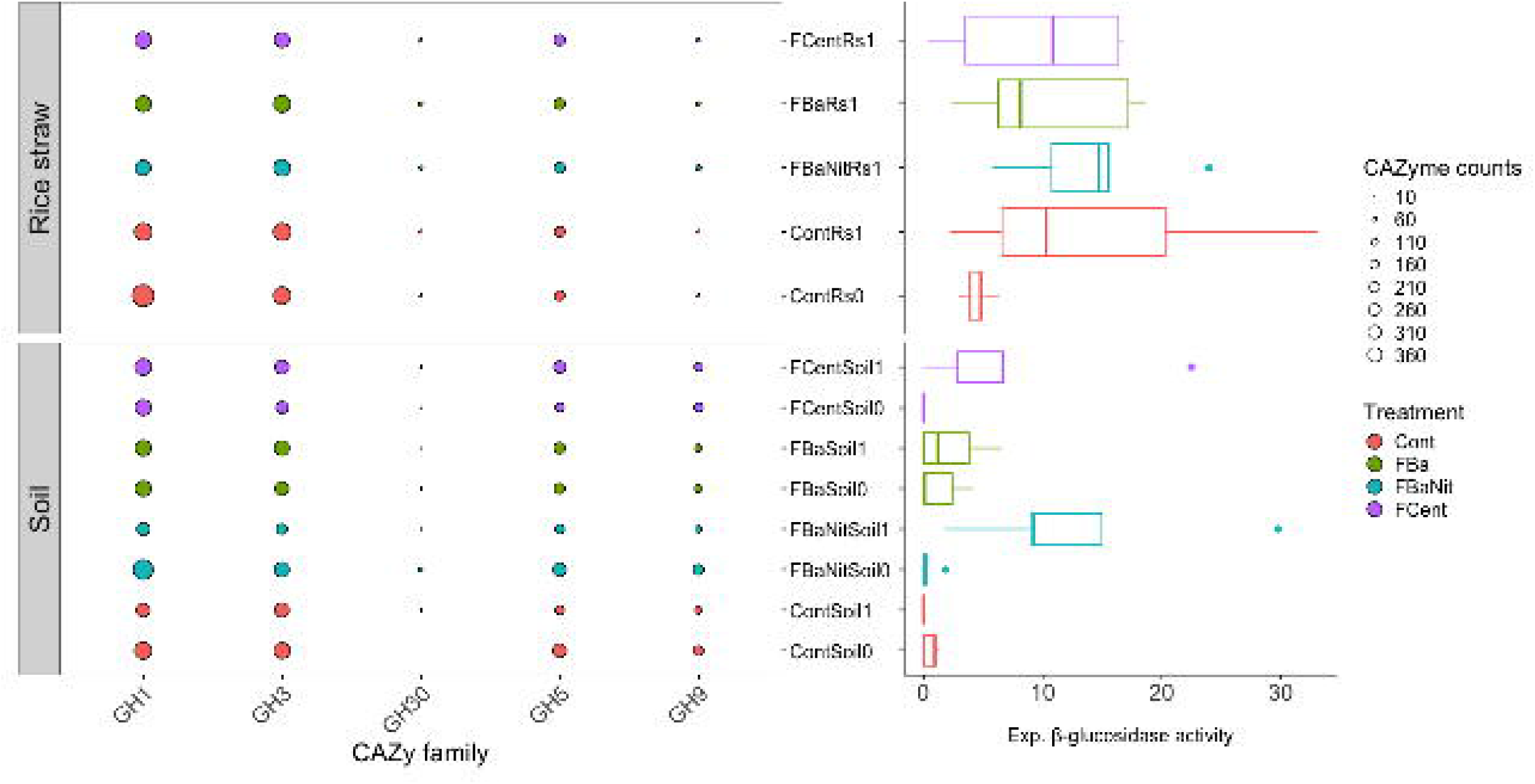
***Left***: bubble plot depicting normalized CAZyme counts for β-glucosidase families (GH1, GH3, GH5 and GH9) per sample; GH glycoside hydrolases. ***Right***: data distribution of experimental β-glucosidase activity per sample. Values displayed on the left are the cumulative sum of each CAZyme family within a determined sample, normalized by the number of coding sequences found in the sample; treatment colors are shared by both the bubble plot and the boxplot.

On the other hand, CAZyme families namely GH1, GH3, GH30, GH5 and GH9 were present within all samples, excepting *Cont* samples at time *0*, where GH30 was absent (***Fig. 8***). Further the contribution from each MAG (before clustering using ANI) to the abundance of specific β-glucosidase families is presented in ***Fig. S16*** and ***Fig. S17*** (***Additional file 1***). In the RS case (***Fig. S16***), it is noticeable how the presence of CAZymes is widespread across several MAGs encompassed by the most abundant phyla Actinomycetota, Pseudomonadota and Bacteroidota, being this former phylum a group with a slight concentration of GH30 modules; special remark must be conferred to *ContRs0_28* as it stands out as a GH9 genes carrier, as well as to *ContRs0_4* (clustered with *FBaRs1_55*) which seems to contain a superior number of GH30.

On the contrary, in soil (***Fig. S17***, ***Additional file 1***), CAZyme β-glucosidases appear to be more localized within the MAGs belonging to the phylum Acidobacteriota, with lower contributions from Actinomycetota. Analogous to the RS situation, a MAG mentioned previously and classified as *Draconibacterirum* (p_Bacteroidota) exhibits exceptional normalized counts of GH30. Nevertheless, in soil matrices at time *0* no experimental activity was detected in spite of the presence of CAZymes related to β-glucosidase activity (***Fig. 8***). This situation prevented the establishment of effective correlations between experimental enzyme activity and the theoretical capacity to metabolize cellulose conferred by the identification of CAZymes in the samples, especially at time *0* in soil. Notwithstanding, we inspected the specific origin of the CAZymes GH1, GH3, GH30, GH5 and GH9 within the treatment that depicted a high β-glucosidase activity (*FBaNit1* in soil) (***Fig. S18***, ***Additional file 1***), and these families, excepting GH30, were found across different taxonomic groups including Acidobacteriota, Actinomycetota, Pseudomonadota, among others; a similar situation was observed in the control sample (*Cont1* in soil) (***Fig. S19***, ***Additional file 1***). The family GH30 mentioned as an exception was only found in MAGs belonging to Acidobacteriota and Verrucomicrobiota; however, this family can be found in other phyla (Bacteroidota and Actinomycetota) within different samples, *FBa* or *Cont* soil samples (***Fig. S17***, ***Additional file 1***).

## Discussion

### Recovery of MAGs with high potential to decompose lignocellulosic material

RS represents a reservoir of both carbon and energy that can be utilized to develop more sustainable agricultural strategies by reincorporating it into the soil. This practice not only would potentially benefit crop cycles, nutrient availability and overall yield, but also it has the power to contribute in the quest to reduce carbon dioxide and greenhouse gases emissions by avoiding the widespread practice of burning the material after each harvest cycle. This work examined the RS decomposition attempted by the application of different biological treatments that include a *Trichoderma* sp. based commercial product, a bacterial strain (*B. altitudinis*) and/or potassium-reducing organic acids.

The MAGs retrieved by this work allowed describing how the microbiota belonging to RS was dominated in at least 50% or above by Actinomycetota and Pseudomonadota, with lower proportions of Bacteroidota, Bacillota and Chloroflexota, whilst, in soil MAGs, Acidobacteriota (the most abundant phylum) governed the community complemented by Actinomycetota and Pseudomonadota, Tectomicrobia and Thermoproteota (Archaea). Interestingly, considering the most abundant phyla in both matrices, only Actinomycetota and Pseudomonadota are shared among them, which suggests communities well-adapted to each specific niche conditions. The presence of Acidobacteriota, Actinomycetota, Bacteroidota, Pseudomonadota, Bacillota and Chloroflexota has been previously documented in Colombian soils at different elevations (between 1000 and 3800 m.a.s.l.) [68]. Likewise, the existence of Actinomycetota and Bacillota has been detected through amplicon sequencing during experiments aimed at establishing a correlation between the microbiota in cacao crops and cadmium mobilization [69].

In terms of MAG taxonomic classification, Actinomycetota (Actinobacteria) was the most prevalent taxonomic annotation, an observation that coincides with reports where this phylum has been recognized by its ability to produce hydrolytic enzyme systems involved in biomass modification [70]. Also, some studies have highlighted that throughout or after rice straw composting Actinobacteria governed the decompounding community, being in some cases almost exclusively the only detected microorganism in the samples [22, 71]. Nonetheless, in RS MAGs the CAZyme distribution within this phylum seems well balanced across the members of the group, whilst in soil MAGs, specific Actinobacteria MAGs (*FBaSoil1_17* and *FBaSoil0_20*) seem to be contributing at an important extent as they contain high number of families related to carbohydrate metabolism namely GH6, GH8, GH26 and GH35.

The relatively high abundance of Pseudomonadota in our experiments constitutes an interesting shared feature by both RS and soil communities since the MAGs belonging to this phylum did not stand out as exceptional carriers of CAZymes. Notwithstanding, Pseudomonadota contains some genera that have been reported with the ability to synthesize thermophilic endo-cellulases, exo-cellulases and β-glucosidases such *Pseudomonas putida* [72], and the genus *Pseudomonas* in general has been referred as part of favorable strategies to biovalorize lignin [73]. Further, after 16S rRNA analysis of colonies isolated from the Keri Lake (Greece), at least one third of the strains were clustered under the Pseudomonadota phylum with a concomitant identification as lignin, xylan or cellulose decomposers. The presence of Pseudomonadota within degradation matrices, however, seems to be dependent on the system as in pile composting samples is not detected [21], whilst it was the most abundant phylum in leaf litter composting systems [74] and in oat straw packed in nylon bags buried in soil containers [75].

Moreover, Acidobacteriota represents the third of the most abundant phylum on the planet, although its practical metabolic capabilities remain as a persistent enigma [76]. This situation relies on the difficulty to cultivate Acidobacteriota species in laboratory environments, and some members of this phylum have been reported to be facultative anaerobic or strictly anaerobic [77]. Consequently, our Acidobacteriota MAGs found in soil can belong to this former class considering that they were remarkably abundant in this matrix, whereas this type of MAGs was not present in RS samples. Further, given its ubiquity, Acidobacteriota species have been found in different environments such as marine sediments, wastewater treatment plants, soil, among others, and they have been recognized to be heavily involved in carbohydrate degrading activity, polyphosphate-accumulating and sulfite/sulfate respiration [28, 78–80]. Nevertheless, despite the fact that Acidobacteriota members have been described as potential decomposers of cellulose and hemicellulose, the experimental enzymatic activity measurements do not agree with the metabolic predictions based on genomic annotations of these microorganisms [77, 78]. This is a concordant situation with our findings as the outstanding carbohydrate-related enzymatic machinery contained within our Acidobacteriota MAGs recovered from soil was not translated into higher rates of RS bioconversion, especially in samples where no β-glucosidase activity was detected.

Specifically in RS mulching systems, Otero-Jimenez et al. (2021) [19] found a significant enhancement in the abundance of Acidobacteria (Acidobacteriota) in the soil after applying RS as mulch for 30 days, and then incorporating it in the soil in a RS management strategy that included application of Fitotripen® and *Bacillus altitudinis,* as we applied in this study. As a result, it seems to confirm that leaving RS as mulch previously inoculated with microorganisms to boost its degradation, may stimulate the increase of Acidobacteriota in the soil, accelerating the degradation of organic matter in this matrix, which in consequence may contribute to lower the soil pH, as in both studies there was a small decrease in pH, that in the case of Otero-Jimenez et al. (2021), was statistically significant.

Furthermore, considering recent works attempting to functionally characterize MAGs retrieved from ongoing or finalized RS-decomposing systems, Santos-Pereira et al. (2023) [29] documented the process of building MAGs with carbohydrate-degrading activity during RS bioconversion in composting systems, where the draft genomes exhibited a high abundance of GH5, GH9 (cellulases) and GH3 (oligosaccharide-degrading) CAZyme families. Nonetheless, these outcomes are not correspondingly aligned with our results since within our both RS and soil MAGs, the most prevalent CAZyme families were AA3 (cellobiose-dehydrogenases), GH1 (oligosaccharide-degrading) and GH10 (endo-hemicellulases); GH3 and GH43 (oligosaccharide-degrading) were remarkably abundant CAZyme within RS MAGs (***Fig. 7***). This situation suggests a theoretical broader spectrum of action of the MAGs hereby described given the variety of the most abundant CAZyme families. However, it is worthy to remark that our results are in accordance with the observations provided by Qu et al. (2023) [22], where their qPCR quantification of defined CAZyme families during pre-inoculated RS composting demonstrated the high expression of GH3, GH10 and GH43 enzymes, precisely some of the most numerous families within our RS MAGs.

Similarly, Kimeklis et al. (2025) [74] recovered 9 MAGs with abnormally high proportion of polysaccharide utilization loci (PULs) relative to their genome size from either straw or leaf litter composting, and within these MAGs, GH43 and GH3 constituted the most abundant CAZyme families, which is consistent with our findings. Although, it is important to mention that Bacteroidota annotation phylum composed more than a half of the selected MAGs by these authors, which in our case was not higher than 20% of the MAGs found in RS samples (***Fig. 2b*** - ***Fig. 3***). Also, from the MAGs retrieved during RS incubation with soil slurries [81], it was possible to establish that the phylum Bacteroidota represents the largest reservoir of GH CAZyme families related to cellulose and hemicellulose degradation mainly in the later stage of the assays. It is noteworthy that Bacteroidota represents the principal phylum in terms of carbohydrate metabolism [82], and this is corresponding with our findings in the sense that more than a third of our RS MAGs with the highest potential to biotransform lignocellulose belong to this phylum (***Fig. 5***).

In respect to the type of enzymes found in higher amounts within the MAGs regardless of the matrix, it was noticeable how cellulose and hemicellulose breaking enzymes, mainly GHs, were the most prevalent CAZymes. Whereas, lignin-associated enzymes, specifically AAs, were found in lower proportions in the MAGs. This situation is related to the fact that cellulose and hemicellulose would be easier to access and hydrolyze to convert it into a more assimilable source of carbon and energy. In contrast, lignin is a complex polymer that requires specialized oxidative enzymes for its biotransformation, and it is metabolized once more labile carbohydrates have been consumed. This sequential degradation pattern has been demonstrated during RS solid state fermentation inoculated with *P. chrysosporium* and *T. viride*, as well as through straw liquid fermentation with a thermophilic composite microbial system [83, 84]. In addition, lignin-degrading enzymes (laccases, manganese peroxidases, and lignin peroxidases) are more commonly found within fungi [85], a scenario supported by the results shown in the section ***Reference-based compositional analysis*** (***Additional file 1***), where RS and soil microbial communities considered in this study were widely dominated by Bacteria, representing this domain around 95% of the annotated sequences.

### Novel uncovered MAGs in Colombian soils

On the other hand, this work allowed the recovery of HQ MAGs with high potential to decompose lignocellulosic materials from both matrices RS and soil whether working together or almost in isolation. In the case of soil MAGs, none of them was taxonomic classified by GTDB-Tk2 at species levels, and in some cases the identification was successful only up to order level. Recent studies carried out in Latin America soils or fresh water have encountered similar situations, where several of the built MAGs appeared to be new species as they could not be identified with the available tools and databases. This is the case of Venturini et al. (2022) [86], who reconstructed 12 MAGs from Amazonian soil, having only two of them with known identity at species level. Also, Rodríguez-Cruz et al. (2024) [87] reported the finding of 325 MAGs (48 Archaea and 277 Bacteria) in the Cuatro Cienegas Basin (Mexico), from which almost half of the Archaea and around 37% of the Bacteria could not be classified even at the genus level. Interestingly, when microbial communities coming from Brazilian sugarcane plantation cultivation soil are grown in liquid enriched medium and then sequenced, an increase in the proportion (more than 95%) of the taxonomically annotated MAGs is obtained as shown by Weiss et al. (2021) [88].

In the specific Colombian case, despite successful efforts to characterize environmental metagenomes in different ecosystems such as bean rhizosphere [89], *Páramo* soils [68, 90, 91], soil from cape gooseberry citrus orchards crops [92, 93], coal mine spoil dump [94], acidic hot spring [95], biocrust [96], among others, using targeted (amplicon) sequencing, to this date there are few endeavors attempting to study local soil-associated microbial communities by recovering and annotating MAGs. This is the case of Herrera-Calderon et al. (2024) [97], who sampled resource islands in a semi-arid zone of the Colombian Caribbean to isolate heavy-metal resistant or tolerant strains; 75 MAGs were recovered from this area, and within these, a high number of resistance genes for Cu, Zn and Ni was detected.

Specifically, considering the abundance and great quality metrics of the MAGs belonging to Acidobacteriota described in this work, as well as their remarkable potential cellulose/hemicellulose decomposers and the lack of proper taxonomic annotation, it is probable that they represent unknown species that can be further characterized through stringent phylogenomics analysis. This bioprospecting task has been subject of active research in recent years as novel Acidobacteriota taxa have been detailed during seafloor sulfur cycling research, and from activated sludge wastewater treatment plants [79, 80]. In a more local scope, recent works have described the presence of Acidobacteriota in South American soils, i.e., Venturini et al. (2022) [86], recovered 41 MAGs from forest and pasture soils in the Amazonian Forest, of which around 12% of them were annotated as members of this phylum.

Similarly, special attention is attracted by the soil MAG classified as *Draconibacterium* (p Bacteroidota) since this 2 Mbp genome concentrates an outstanding number of GH CAZymes, covering a wide range of activity in terms of cellulose and hemicellulose biotransformation. This finding suggests that even though Bacteroidota presence in the soil matrix is scarce, it seems that a single taxon (or a few taxa in the case of RS samples) has a major role in carbohydrates metabolism. *Draconibacterium* species have been mainly isolated from marine sediments, and at the moment of writing this report, they have not exhibited special abilities to metabolize cellulose [98–103].

### β-Glucosidase experimental activity in RS and soil samples

The experimental enzymatic validation carried out in this work did not allow establishing a determined effect conferred by *Trichoderma* sp. nor *Bacillus altitudinis* IBUN 2717 alone even though multiple studies have underlined the benefits this kind of consortium application brings to lignocellulose-decomposing systems [10, 12, 14, 20]. However, increments in β-Glucosidase activity were statistically significant only for the treatment *FBaNit* in soil, where this treatment could significantly enhance the enzymatic activity compared to *Cont* samples. As a result, the addition of an inorganic source of nitrogen had an effect on the gene expression related to carbohydrate metabolism rate rather than on the microbial composition or presence of higher numbers of genes encoding CAZymes.

Similar to this work, strategies that include the management of carbon to nitrogen ratio (C/N) in combination with microbial inoculants have demonstrated to be a successful strategy for the degradation of RS [5, 104]. For instance, Cruz-Ramírez et al. (2017) [18] found the application of urea, as well as the same microbial consortia used in this work, during RS decomposition to be advantageous for plant growth after RS incorporation in greenhouse experiments. Analogously, Lio et al. (2023) [104] enhanced wheat degradation that led to incremented rice growth and yield in about 13% within a rice-wheat rotation system by applying ammonium bicarbonate, plus straw-decomposing microbial inoculants (SDMI). Moreover, using a high amount of urea per surface area (97.5 kg.ha^1^), Kalkhajeh et al. (2021) [5] reached an increase up to 13.6% in grain yield of rice in a paddy soil when the nitrogen was co-administered with SDMI. These improvements on lignocellulose breakdown given by the addition of additional nitrogen sources are given by its influence on enabling an enriched amino acid anabolism, and hence a boosted protein synthesis, including enzymes [105]. In consequence, extra nitrogen during straw decomposition systems promotes microbial growth and reproduction, thereby elevating the metabolic activity within the microbial community [5]. Not only external nitrogen application can affect the microbiota directly, but also it can directly alter other conditions related to microbial activity and enzyme releasing such as the pH; data recently published depicted a notable acidification after addition of readily available nutrients during lignocellulose degradation in paddy soil [106].

On the contrary, our attempt to correlate the presence of GH1, GH3, GH30, GH5 and GH9 CAZyme families with β-glucosidase experimental activity was not consistent. For instance, the normalized counts of four out of the five families were decreased after 30 days in *FBaNit* treatment in soil, even when an increment in the experimental activity was observed. Additionally, considering that *FBaNit* treatment promoted a higher β-glucosidase activity, no significant differences were found in terms of number of CAZyme gene copies compared to the other treatments. This scenario indicates that the limiting factor would not be the an increased number of genes directly associated with β-glucosidase but the expression and/or specialization of those genes,

Finally, even though we could detect the experimental effectiveness of the treatments by quantifying the bioconversion given by β-glucosidases, the fact that no experimental β-glucosidase activity was observed even when the presence of specific CAZymes was established underlines the necessity to determine direct correlations between CAZyme abundance and experimental activity in lignocellulose-based degrading systems. To achieve this purpose multiple approaches have been reported encompassing fosmid library construction, qPCR for specific CAZyme families or the use of microarrays [22, 29, 107]. These correlations can be supported by identifying microorganisms, or communities, with superior enzymatic activity through specific methodologies as, for instance, microplate-based screening (Biolog MT2) [108].

## Conclusions

This study highlights the potential of leveraging biological strategies to enhance rice straw (RS) degradation through the application of biological strategies. The co-application of inorganic nitrogen to balance the C:N ratio and a microbial consortium resulted in increased βIi.Glucosidase activity and variations in the proportions of genes encoding specific enzyme families in soil samples, which may be related to the observed differences in enzyme activities.

MAG reconstruction and mining of carbohydrateIi.active enzymes enabled the recognition of probable previously unknown microorganisms with remarkable capabilities to degrade RS, i.e., MAGs recovered from RS samples that were clustered under Bacteroidota phylum, albeit without proper assignment of a species name. Similar situation was documented for soil MAGs, in which none of them was fully taxonomically annotated based on a phylogenetic tree that contains more than 580,000 genomes (GTDB). Moreover, many of the MQ or HQ MAGs belonging to the phylum Acidobacteriota demonstrated to account for great lignocellulose-degrading potential as they enclose a broad diversity of CAZymes related to the breakdown of agricultural subproducts, including enzyme families highly reported to be involved in RS bioconversion. Among several great candidates from soil MAGs to lead RS decomposition, a MAG annotated as *Draconibacterium* constitutes an interesting case of study that requires further analysis as it is not known to perform metabolic tasks related to lignocellulose metabolism.

Our work serves as a step forward in the elucidation of alternative biocatalysts methods that lead to enhanced lignocellulosic biomass bioconversion for various industrial applications, and it contributed to unveil the Colombian soil and biological material unexplored microbial diversity. Admittedly, the challenge remains in studying the expression levels of the carbohydrate-degrading enzymes within the genomes, as well as in the need to activate the catalytic enzymes by enhancing the proposed bioconversion strategies.

## Declarations

### Ethics approval and consent to participate

Not applicable.

### Consent for publication

There is no conflict to consent for publication.

### Availability of data and materials

The raw sequences can be found under the BioProject <u>PRJNA1222139</u>. HQ and MQ MAGs, low-quality-bins, non-representative MAGs and supplementary files can be accessed on <u>Zenodo</u> <u>(10.5281/zenodo.14882745)</u>; mock community information and metadata can be retrieved from the same Zenodo repository. Also, with aim of easing reproducibility of the results hereby presented, scripts used to run the different tools are available at the same repository, as well as software outputs (dbCAN3, Prokka, COGClassifier, FastANI, MAGFlow, FastQC, nf-core/mag, FetchMG, GTDB-Tk2 and Kraken/Bracken) and knitr/executable R and Python downstream-analysis scripts. A supplementary and interactive view of the figures can be reached on the following Github page (https://jeffe107.github.io/metagenomics-report/), and an online instance of BIgMAG showing quality metrics and taxonomic annotation of the clustered MAGs is accessible at https://bigmag.online/.

### Contributions

JYG performed reference-based taxonomic profiling, MAG assembly, annotation and curation, manuscript writing, and data integration, visualization and deposition. NNM carried out field experiments, DNA extraction, physico-chemical measurements, enzymatic activity quantification, quality control checks and manuscript writing. VOJ contributed to manuscript writing, reviewing and editing. DUV, EBH and LF supervised and oversaw experiment development and data analysis. All authors participating during the study framework conceptualizing, conceiving the overall work and manuscript preparation.

## Acknowledgments

The authors acknowledge the assistance received by Dr. Maria Fernanda Alvarez, Marcela Pineda and Cristian Herrera from The Rice Program at the Alliance of Biodiversity International and International Center for Tropical Agriculture (CIAT), who contributed to this work at allowing experimenting in their rice fields and helping during the assay execution. JYG specially thanks the Federal Commission for Scholarships for Foreign Students (FCS) for their support through the Swiss Government Excellence Scholarship.

## Additional files

***Additional file 1***. **Supplementary Results and Figures** (Suppl_results_GRMAoRSDEUMAGwHPtDLR.docx). Microsoft Word document with supplementary figures and complementary results description and discussion.

***Additional file 2***. **Supplementary Tables** (Suppl_tables_GRMAoRSDEUMAGwHPtDLR.xlsx). Microsoft Excel workbook with supplementary tables.

