## Additional file 1 for "Genome-Resolved Metagenomics Analysis of Rice Straw Degradation Experiments Unveils MAGs with High Potential to Decompose Lignocellulosic Residues"

**^e^**Bioinformatics group, Biotechnology Institute, National University of Colombia, A.A 14-490, Bogotá D.C., Colombia

**Abstract**

**__________________________________________________________________________________**

**Background**

Rice is one of the top three crops that contribute 60% of the calories consumed by humans worldwide. Nonetheless, extensive rice harvesting yields more than 800 million tons of rice straw (RS) per year globally, generating a byproduct that is often difficult for farmers to manage efficiently without burning it. As a result, millions of tons of carbon dioxide and greenhouse gases are released, causing issues such as respiratory problems, soil degradation, and global warming. In this work, we explore the biological decomposition of RS through the application of microbial consortia from a metagenomics perspective.

**Results**

We applied different treatments to RS placed in a mulching setup during experiments carried out in Colombian rice fields, using various combinations of a *Trichoderma*-based commercial product, the bacterial strain *Bacillus altitudinis* IBUN2717, inorganic nitrogen, and a mixture of potassium-reducing organic acids. Before inoculation and after 30 days of treatment, we characterized the microbial community on the RS surface and from the bulk soil by performing a reference-based compositional analysis, and reconstructing and functionally annotating Metagenome-Assembled Genomes (MAGs). High-quality MAGs with great potential to decompose RS, represented by the extensive number of carbohydrate-active enzymes, were recovered. Soil MAGs taxonomic classification indicates that they may represent potential novel microbial taxa. At the same time, the main part of the RS MAGs with superior lignocellulose-degrading capacity were affiliated under Actinomycetota and Bacteroidota phyla. Moreover, β-glucosidase activity measurements indicated an increased RS degradation after the application of the treatment that included inorganic nitrogen.

**Conclusions**

This contribution underscores the possibility of promoting RS degradation through the application of biological strategies. Further, the newly unveiled MAGs with high RS-degrading potential provide a valuable resource for exploring the functional potential of previously uncharacterized microbial diversity in Colombian agricultural ecosystems, including microorganisms that have not been previously reported as remarkable lignocellulose decomposers.

**Supplementary Results and Figures**

**Area of study**

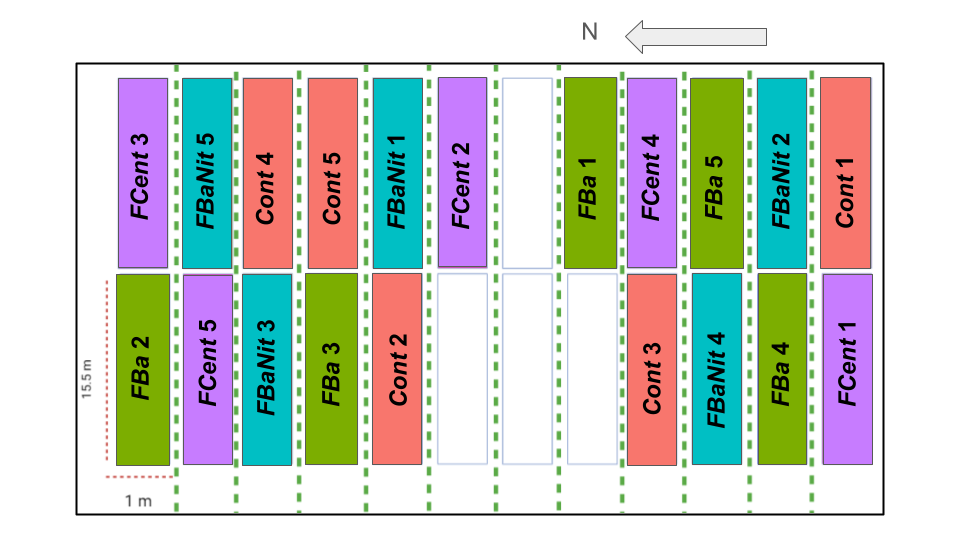

***Fig. S1.*** Field diagram of plots and yard distribution, illustrating yard sizing and color plotting distribution on the field. Green dot lines indicate rice furrows without treatment used as a natural barrier between plot treatments. Blank spots represent spaces without treatments due to field irregular terrain impeding a treatment application. Arrow indicates magnetic north.

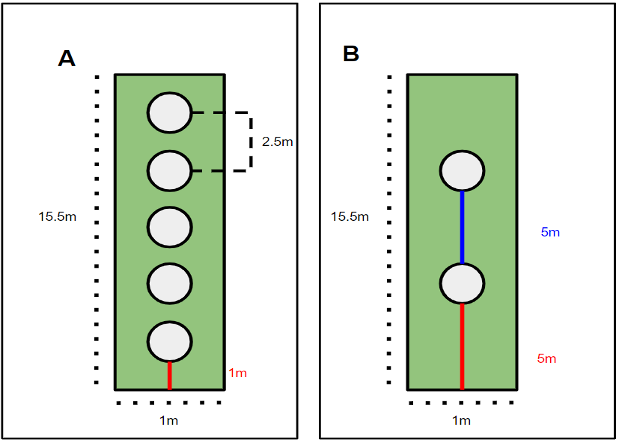
**Field experiments**

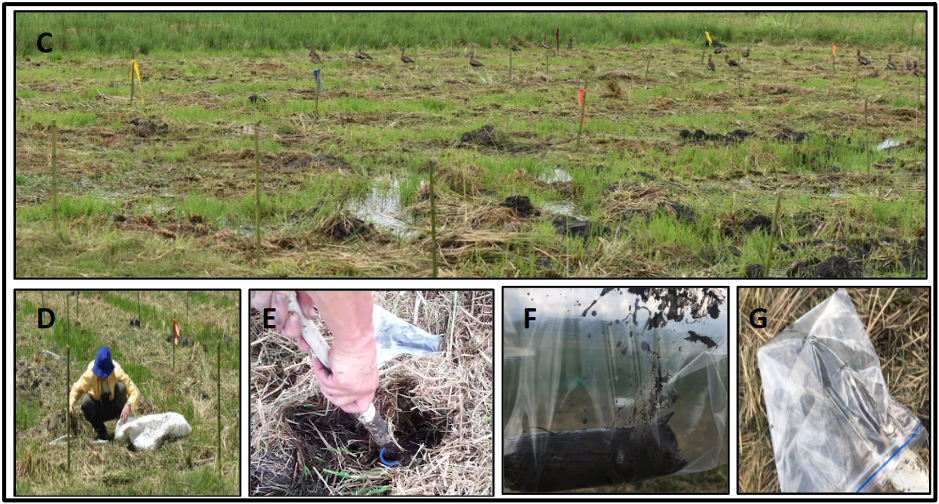

***Fig. S2***. Illustration of sampling points for soil and straw for physicochemical and enzymatic analysis on field. ***A*** distribution of soil sampling points within each plot; ***B*** distribution of rice straw sampling points within each plot; ***C*** field picture showing the marks used for the replicas and color code per treatment: light pink represents *Cont* plots, blue *FBa* plots, yellow *FBaNit* plots, and violet *FCent* plots; ***D*** RS manual collection; ***E*** tube probe for soil sampling; and ***F***, ***G*** soil sample disposal in marked Ziploc® bags.

**Soil physicochemical analyses**

In order to determine the effect of the treatments under field conditions, soil physicochemical parameters were evaluated including pH, total soil carbon, total soil nitrogen and available phosphorus after 30 days of trial. These parameters demonstrated changes over time, where all treatments positively affected carbon sequestration (***Fig. S3***), increasing total soil carbon after 30 days; however, these increments were statistically insignificant, according to the Analysis of Variance (ANOVA) (***Table S1***, ***Additional file 2***). The soil C/N ratio demonstrated temporal variation in response to some of the treatments. According to the measurements, at time *0* in the *Cont* plots, the C/N ratio was 16:1, and one month later, this ratio dropped to 10:1; an analogous scenario was observed for *FBaNit* and *FCent* treatments, where it descended until 12:1 and 11:1, respectively. In the case of *FBa* treatment the C/N ratio remained at the same initial value 1:16. Similarly, within the 30-day timeframe, soil-available phosphorus decreased in all cases, while pH prevailed around the same value, 7.69.

**Enzymatic activity**

***Fig. S4*** shows the measurements of different enzyme activities including protease, acid, and alkaline phosphatase. In regards to protease activity in RS samples, these matrices exhibited a prominent reduction in protease activity after 30 days of treatment application in all cases. Whilst, in soil samples, C*ont* treatment did not show distinguishable activity after 1 month, while *FBa* and *FBaNit* seemed to decrease protein catalysis within the same range of time. In one of the replicates belonging to *FCent*, this enzyme activity was increased, although across treatments the coefficient of variation was high, and hence it is not possible to determine a clear effect of the treatments related to this enzyme in soil matrices. Nonetheless, no significant differences were detected among treatments by the Kruskal-Wallis test.

In the case of acid phosphatase, all RS samples depicted an absence of activity both at time *0* and time *1*. Whilst, in soil samples, this enzyme showed a variable behaviour, where in *Cont* an increase of the activity was measured at time *1*, as well as in *FBa* samples; *FBaNit* treatment did not appear to have any effect in soil matrices. However, in RS samples, the alkaline phosphatase activity was augmented notably in *FBaNit* and *FCent* treatments, while the detection on *Cont* samples was moderately enhanced after 30 days; in soil matrices, the quantification of this enzyme did not reveal any treatment effect since no changes were observed after 1 month of assays. Again, the variability of these enzymatic measurements was remarkably high, and therefore general trends or significant differences were not possible to establish.

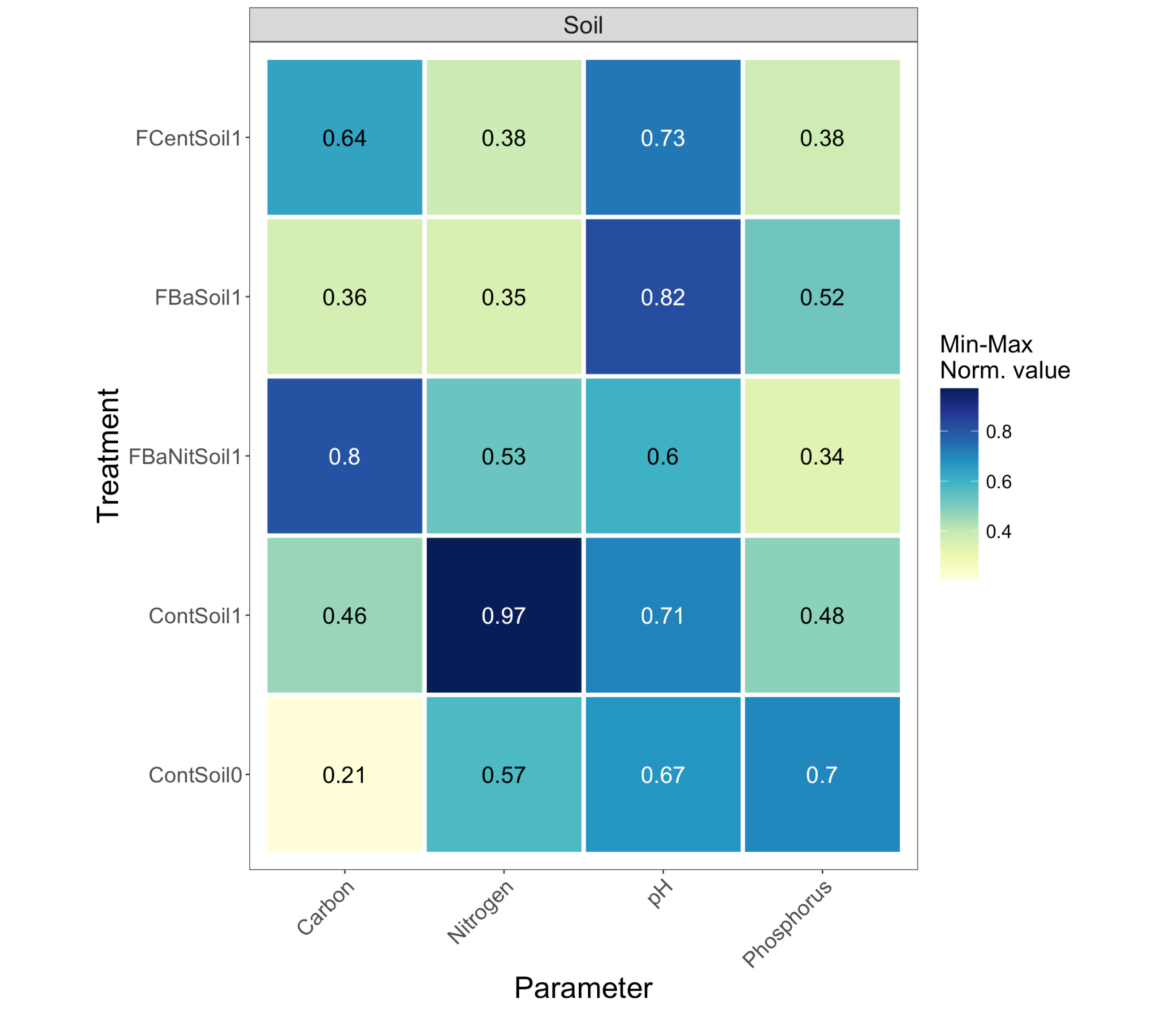

***Fig. S3***. Experimental measures of physicochemical parameters in **soil** samples including total carbon, total nitrogen, pH and phosphorus. The data were normalized using the Min-Max method to adjust the raw values to the same scale; the displayed numbers represent the mean of the quantified parameter per sample. Analysis of Variance (ANOVA) evaluating the effect of treatments on each variable are presented on ***Table S1***, ***Additional file 2***.

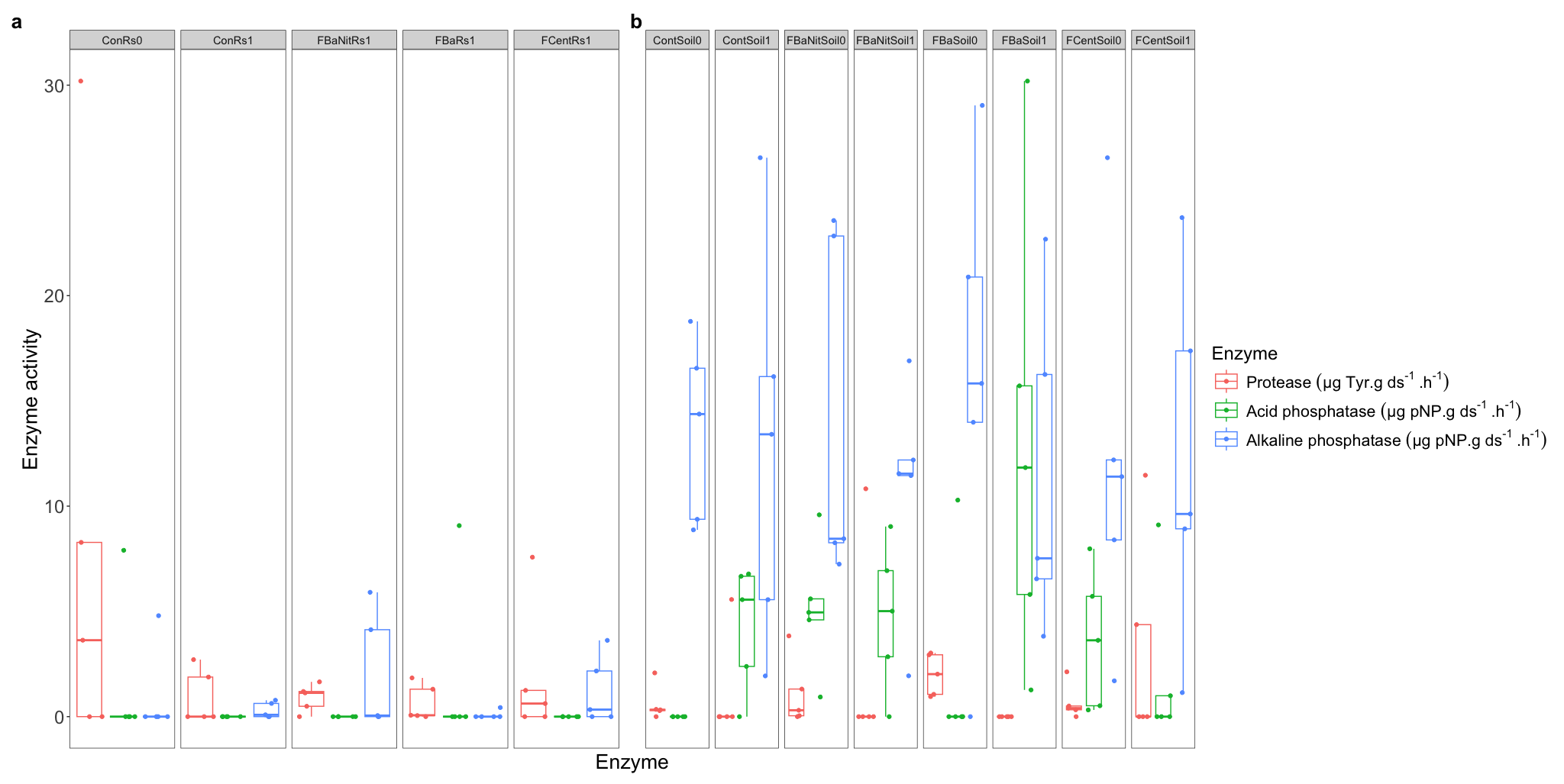

***Fig. S4***. Experimental protease, acid, and alkaline phosphatase activities of **RS** (***a***) and **soil** (***b***) samples. The enzymatic activity was measured according to the guidelines established by Alef & Nannipieri (1995).

**Reference-based compositional analysis**

In our work, we carried out different experiments applying different biologically-based treatments over mulched RS as an attempt to both comprehensively study the microbial community and enhance the degradation rate. Firstly, the integrity of the methodology to extract and sequence the nucleic material was validated by inclusion of a mock community as an extra sample. The abundance values of each community member (***Fig. S5***), annotated by Kraken2 and re-estimated by Bracken, is in agreement to the composition stated by the provider, corroborating the well performance in general of the chosen DNA isolation and sequencing methods.

The taxonomic classification and abundance quantifying of the sample sequences demonstrated that both environments, RS and soil, are primarily governed by the domain Bacteria, since more than 95% of the annotated sequences belonged to such domain (***Fig. S6a***). Archaea, Eukaryota, and Viruses were detected in low proportions, with no pronounced differences in the relative composition of these domains between the two matrices; soil matrices exhibited a higher diversity in domain composition compared to RS.

Moreover, α-diversity metrics of prokaryotic species, measured by Shannon and Inverse Simpson indices (***Fig. S6b***), exhibited a higher diversity in soil than in RS across all treatments. Within each matrix, the metrics were consistent among treatments (*FBa*, *FBaNit* and *Fcent*), including the *Cont* samples, with a few exceptions of replicates that show deviated values. In addition, β-diversity, estimated using PCoA based on Bray-Curtis distances (***Fig. S6c***) and UMAP on Aitchison distance (***Fig. S6d***), revealed distinct clustering of microbial communities by matrix type (RS vs. soil), indicating significant compositional differences. On the other hand, within each matrix, treatment groups overlapped considerably, suggesting that treatments had minimal impact on the general community composition. Being so, the depicted clusters by both methodologies suggest that the matrix type is the primary determinant of microbial community structure. This former statement is supported as well by the fact that on average 65% of the RS reads were annotated by Kraken2 to at least the root taxon, while only 35% of soil reads were classified to this category (***Fig. S9***). This scenario highlights the higher difficulty for the software to assign taxonomic ranks in soil samples given its intrinsic complexity and the limited scope of even the larger public available database. Also, it confirms the high diversity of the microbial communities found in Colombian bulk soil, being remarkably greater than the diversity values depicted by the RS surface by this study.

Furthermore, the compositional variability of the samples (***Fig. S6e***), measured as distance to the centroid in a multivariate space computed with the Aitchison distance among samples, seemed to be greater for RS samples compared to soil regardless of treatment or time point. While the treatments induced some dispersion in the data, mainly in RS samples, their impact was modest compared to the inherent variability between matrices. This situation gives insights for an observed stability of the soil microbial community across time and treatment without subtracting importance to the complex and heterogeneous nature of the soil. Whilst in the case of RS samples, although a higher compositional variability is displayed, this is the result of the effect of time on microbial composition rather than the effect of treatments, as noted below. The ANOVA test on the average distance showed no significant differences in terms of homogeneity of group dispersions (variances) among samples (***Table S2***, ***Additional file 2***).

Going further to evaluate if significant differences in microbial community composition explained by Time and Treatment factors can be found, multiple PERMANOVA analyses were performed with several distance/dissimilarity metrics (Aitchison, Bray-Curtis, Euclidean and Jaccard) (***Fig. S6f***). In the case of RS samples, Treatment accounted for a larger proportion of the variance; however, the strongest effect on the microbial community shape was caused by the factor Time (*p* < 0.05), reflecting the effect of temporal changes; this situation could be likely due to microbial turnover and/or substrate degradation, although these hypotheses require in-depth investigation. Meanwhile, in soil samples, the factor Treatment accounts for a greater effect size as compared to the portion of the variance explained by Time effects, without pronounced differences among treatments regarding the impact on the community composition. In this sense, *p*-values suggest that the microbial configuration does not significantly differ among samples belonging to the same matrix (*p* < 0.05). Note that the outcome values were consistent regardless of the input matrix for the test.

In terms of the microbiota distribution, the dominant bacterial phyla in both matrices were Pseudomonadota and Actinomycetota, with Bacillota, Bacteroidota, Cyanobacteriota, Myxococcota, and others contributing to a lesser extent (***Fig. S7a***). In RS samples, Bacillota exhibited higher relative abundance in the *Cont* samples at the beginning of the experiments with a pronounced drop at time *1* in all treatments, while Acidobacteriota was more prevalent in soil samples compared to RS matrices.

Additionally, ANCOM-BC2 test at genus level outcomes corroborates PERMANOVA findings by demonstrating the strong influence by the factor Time on the microbial community composition in RS samples causing sharp drops in some specific genera (***Fig. S6g***). Specifically, the genera *Lactococcus* and *Aeromonas* were heavily impacted after 30 days in all of the treatments, including *Cont* samples, while *Priestia* was affected across treatments, except for *FCent* samples. In *FBa* matrices *Acinetobacter*, *Paenibacillus* and *Morganella* were the most depleted genera at the end of the experiments; as well as *Deftia*, a feature *FBa* samples shared with *Cont* matrices. *Exiguobacterium* was diminished only in *FCent* replicates. Interestingly, when switching the control group against which ANCOM-BC2 performs the comparisons to *Cont* samples at time *1*, no discrepancies are detected among treatments, highlighting a weak power of the treatments to induce changes over the microbiota by themselves, and outlining that the observed changes were almost exclusively generated by the time elapsed during the experiments. This situation may rely on reasons such as high replicate variability within treatments in specific taxa, or microbial resilience over time with some species taking over the community, leading to convergence between treatments and control samples; as a result, insubstantial effects might not have passed the significance threshold given the stringency with which ANCOM-BC2 controls for false discovery rates (FDR). Nevertheless, this is precisely the strength ANCOM-BC2 accounts with compared to other similar methodologies that would show significant results without considering the compositionality of the microbiome data and the intrinsic sampling fraction that must be corrected before determining differentially abundant taxa.

Remarkably, in RS matrices at species level, *B. altitudinis* was detected by Kraken2 transversely in all treatments, including *Cont* and *FCent*, in which this strain was not inoculated. This indicates a native nature of *B. altitudinis* within the environments considered in the present study. Nonetheless, *B. altitudinis* abundance variability among replicates from the same samples was high, and in the case of this genus, metagenomic classifiers may struggle to distinguish among them since members of this genus can be exceptionally similar in genomic terms. On the contrary, at time point *1*, the soil community seemed wholly resilient given the uniformity of the abundance distribution across treatments. A fact supported by ANCOM-BC2 results, considering treatment plus time as the fixed formula, where not a single significant differentially abundant group at genus level was reported by this methodology. Notwithstanding, these observations contrast with the fact that more than 50% of the annotated genera are present in less than 3% of the total microbiota (***Fig. S7b***, ***Fig. S8***).

Although eukaryotic organisms only represent around 3% of the total annotated sequences in both RS and soil, it is worthy to note the abundance expansion of the phylum Ascomycota at time *1* in RS samples, being this phylum more abundant in RS than in soil samples in any case (***Fig. S7c***). However, it is particularly noteworthy that *Trichoderma* sp. does not appear as one of the dominant genera in none of the samples after 30 days of treatment evolution (***Fig. S7d***); it was detected in negligible proportions (< 0.01%), and the reported species by Kraken2 were *T. breve*, *T. asperellum* and *T. atroviride*, where none of them was part of the commercial product applied (Fitotripen®). This fact would suggest that *Trichoderma* activity is started shortly after inoculation, and then it transitions back to its sporulated state, for which the DNA extraction protocol followed in this study is not suitable.

**Differences among matrices and treatment effects from read-based taxonomic profiling**

In principle both matrices, RS and soil, are governed almost exclusively by Bacteria, leaving only a 5% of the environment to Eukarya, Archaea and Viruses. Consequently, the microorganisms of this life domain carry the burden of metabolizing carbon from organic sources. This scenario is analogous to the ones described during the thermophilic stage of compost samples by Santos-Pereira et al. (2023) [[1]](https://www.zotero.org/google-docs/?GnISyg), and it can be found across different environments such as limestone aquifer, where Bacteria composes between 94% to 98% of the microbial community [[2]](https://www.zotero.org/google-docs/?zLTyVX).

From the diversity analysis, both α and β, it is possible unequivocally to establish differences between soil and RS samples, where soil matrices exhibited a higher diversity and complexity; an unsurprising situation given the vast number of academic works stating such an incredibly high abundance of species in soil environments [[3, 4]](https://www.zotero.org/google-docs/?BpqMAD). In addition, the microbiota associated with this type of matrices has been proven to be clearly different, and dynamic, during the aerobic and anaerobic incubation of RS and soil [[5]](https://www.zotero.org/google-docs/?ej1RON).

Furthermore, a following of the evolution of the field plots depicted how the soil samples remained steadily stable throughout time showcasing a resistant and resilient microbial community as established by Allison & Martiny (2008) [[6]](https://www.zotero.org/google-docs/?PeRdYc). In the meantime, after 30 days of treatment application, the taxonomic profiling showed a sharp reduction in the abundance of the Bacillota phylum on the RS surface. Among the most abundant genera, Acinetobacter, a member of this phylum, exhibited a proportional abundance diminution. This is particularly intriguing mainly for *FBa* and *FBaNit* treatments, in which a member of this phylum, *B. altitudinis*, was extensively included at the beginning of the experiments. Recently, several studies have identified the contraction of the Bacillota phylum abundance during lignocellulose decomposition in different environments. For instance, Sarma et al. (2022) [[7]](https://www.zotero.org/google-docs/?rZXLK9) found a reduction of Bacillota (formerly Firmicutes) in 10 days after the initiation of RS pile composting, Kimeklis et al. (2025) [[8]](https://www.zotero.org/google-docs/?p9avJb) detected a drop in this same phylum after 111 days of leaf litter composting, as well as Sun et al. (2023) [[9]](https://www.zotero.org/google-docs/?ABibZG), who reported a significant drop in Bacillota abundance after 60 days of wheat straw decomposition packed in nylon bags and embedded into the soil. This observed shrinkage on Bacillota abundance may be related with an increase in the temperature of the fields where the experiments were carried out given that authors as Sarma et al. (2022) [[7]](https://www.zotero.org/google-docs/?UI5AFT) have attributed this behavior to growth inhibition at temperatures above 40°C reached by the compost piles after 10 days of assay (early thermophilic stage).

Captivatingly, *Trichoderma* sp. detection across all treatments in both matrices, at time *0* and *1*, was almost negligible posing puzzling questions in regard to the activation, growing and development of this important component of the treatments. Nonetheless, assays carried out by Organo et al. (2022) [[10]](https://www.zotero.org/google-docs/?fsv70q), where degradation of RS was performed through the litter bag technique, and *Trichoderma* was used as activator, depicted a rapid reduction in the CFU counts of *Trichoderma* after 14 days when the bags were placed on soil surface, and being almost undetectable after 42 days. Consequently, given the timeframe and the superficial application of *Trichoderma* considered in the present work, probably the spores of this fungus contained in the commercial product Fitotripen® germinated speedily, grew abundantly, and by the time the plots were sampled (after 30 days) its abundance had already declined enough to making it insignificant to be quantified by our pipeline.

Finally, through the reference-based compositional analysis of RS before and after treatment application, it was possible to establish how the RS microbial community differs from soil microbiota to a large extent in both α and β-diversity in spite of the physical closeness of these microenvironments. Also, the soil stability and resilience were established since its microbial population remained the same during the time window of this work, and in both matrices the microbiota was governed by Bacteria given its abundance above 95%. Within this Bacteria, Actinomycetota, commonly related to lignocellulose decomposition, represented the highest portion of the microbial population, and Bacteroidota, the principal phylum in terms of carbohydrate metabolism according to different reports, resulted as a secondary member of the community according to this read-based analysis. Meanwhile, in soil samples, Acidobacteriota, a phylum recognized by its bioconversion activity, was prevalent across treatments; however, significant differences among treatments in terms of community composition were not detected in field trials.

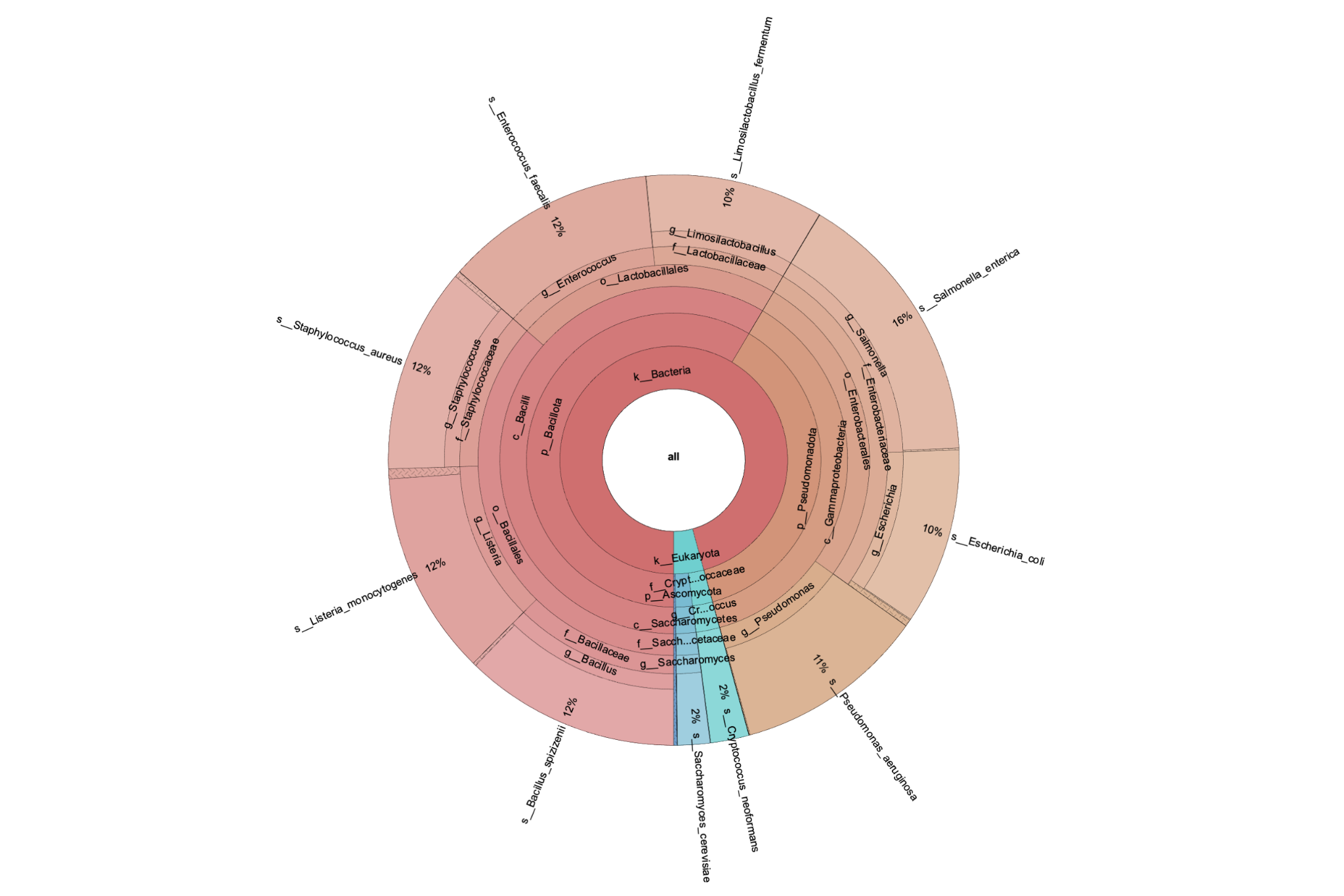

***Fig. S5***. Screenshot of a Krona plot displaying the taxonomic classification and relative abundance, performed with Kraken2 coupled to Bracken, of the reads belonging to the mock community sequenced as part of quality control for the DNA extraction process.

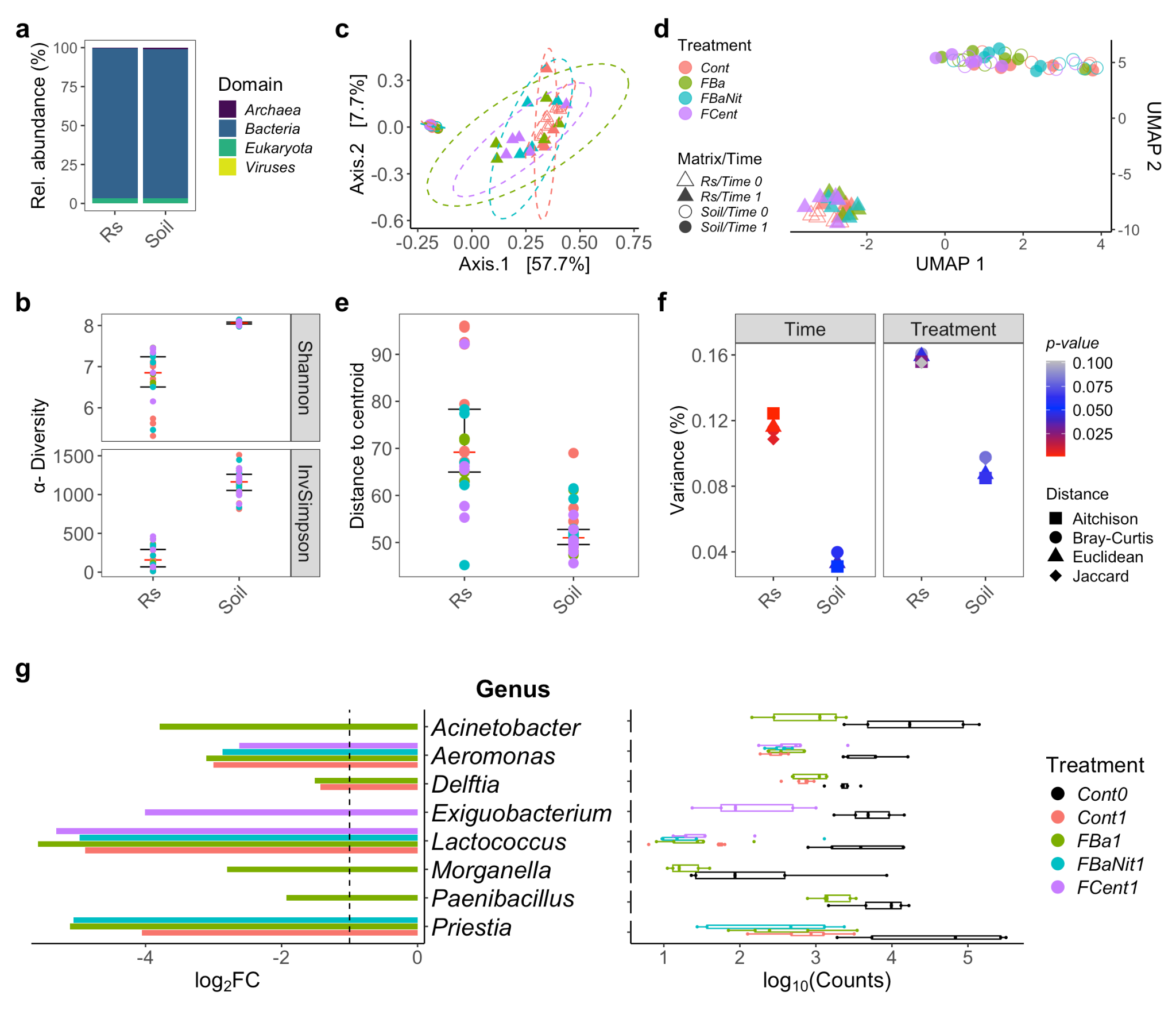

***Fig. S6***. Taxonomic profiling of the sequences using Kraken2 coupled to Bracken: ***a*** Relative domain abundance per matrix source, merging all the replicates per sample. ***b*** α-Diversity, colored by treatment, measured through Shannon and Inverse Simpson indices. ***c*** Sample ordination through Principal Coordinate Analysis (PCoA) using a Bray-Curtis dissimilarity matrix as input. The ellipses represent 95% confidence regions around the centroids of each group calculated by the assumption of multivariate t-distribution. ***d*** Two-dimensional Uniform Manifold Approximation and Projection (UMAP) embedding of β-Diversity estimation of the samples based on the Aitchison distance. ***e*** Compositional variability of the samples, measured as distance to the centroid in a multivariate space computed with the Aitchison distance among samples. ANOVA analysis results show no significant differences in terms of group dispersion. ***f*** Proportion of the variance explained by the factors Time and Treatment, segmented by matrix source, in Permutational Analysis of Variance (PERMANOVA) implemented according to different distance/dissimilarity metrics; the significance of each factor is determined by the symbol color. ***g*** Selected differentially abundant genera in RS samples based on Analysis of Compositions of Microbiomes with Bias Correction 2 (ANCOM-BC2). ***Left***: log_2_FC of treatments to *Cont0*, and ***Right***: boxplots of selected genera abundance (counts) in log_10_ scale. All displayed genera exhibit *p*-value < 0.05 adjusted with Holm–Bonferroni method for multiple comparisons. *Cont0* and *Cont1* in RS comparisons indicate *Cont* samples at time *0* and *1*, respectively. Differences throughout time for all treatments against *Cont* samples at time *0* are shown. All the analysis from ***b*** to ***f*** were carried out only with Bacteria data and normalized counts as Counts per million (CPM). The treatment colors are transversal to plots ***b***, ***c***, ***d***, ***e***, ***f*** and ***g***. ANCOM-BC2 was implemented with agglomerated data at genus level, and with the most abundant taxa (> 0.5%).

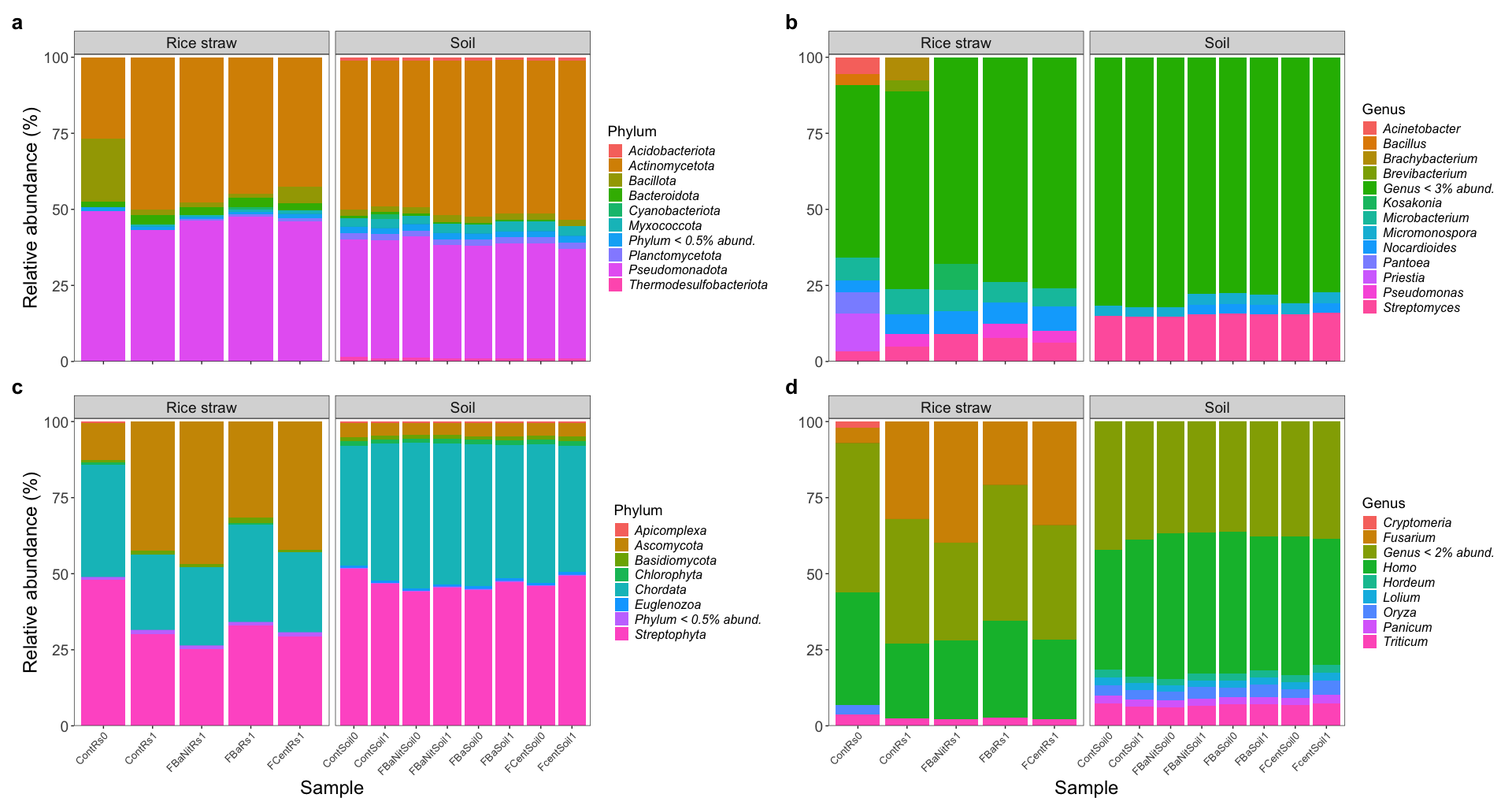

***Fig. S7***. Relative **phylum** (***a***, ***c***) and **genus** (***b***, ***d***) abundance of agglomerated data (**Bacteria**, ***top***; **Eukaryota**, ***bottom***). Displaying abundance values that belong to the portion of the reads annotated in each domain (*see* ***Fig. S6a*** and ***Fig. S9***). Replicates were merged, and the panels are split by matrix source. On phylum plots, only phyla with relative abundance above 0.5% are displayed, whilst on genus plots genera above 3% and 2% are depicted for Bacteria and Eukaryota, respectively.

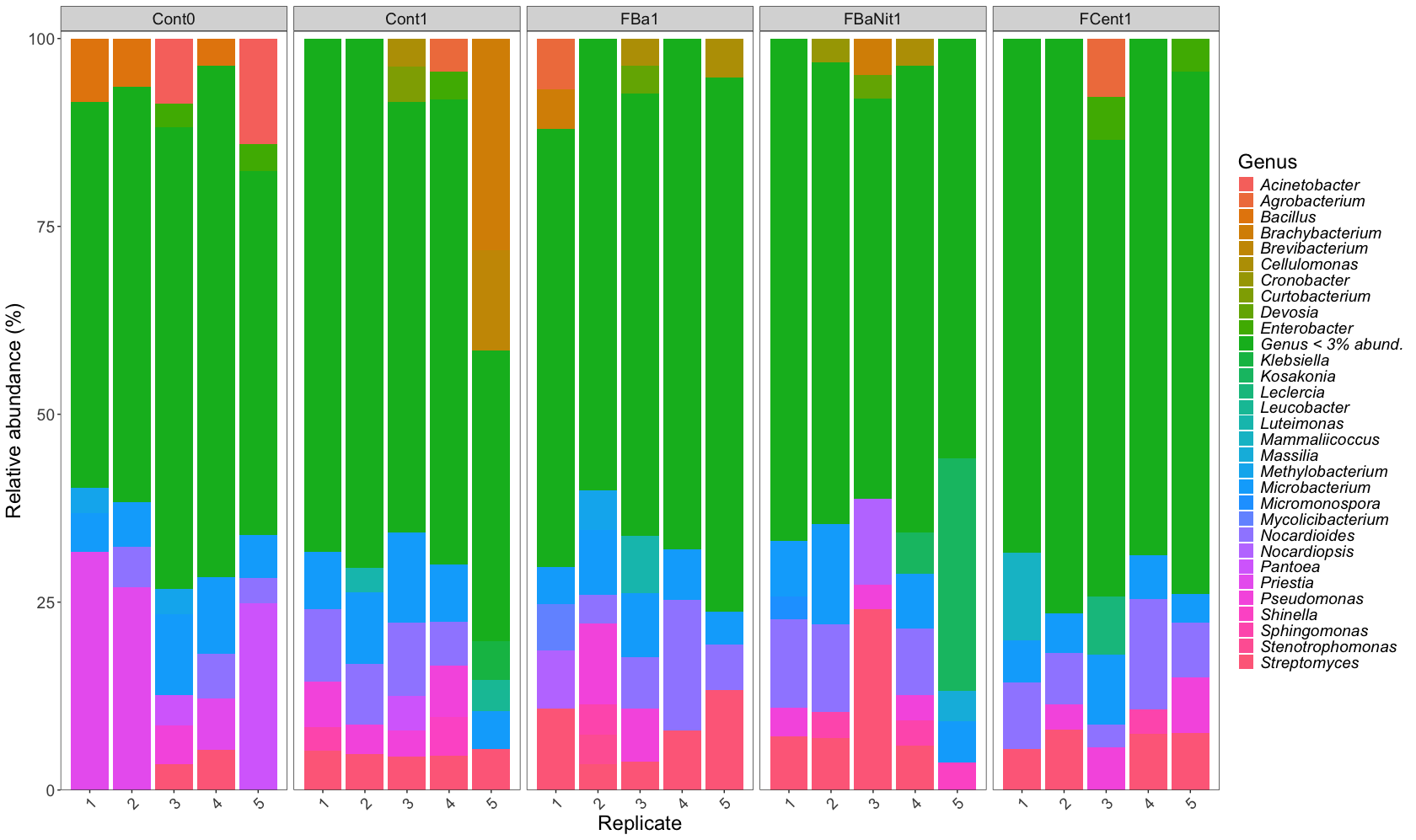

***Fig. S8.*** Relative genus abundance per replicate of agglomerated **RS** data with panels split by treatment. Only genera with relative abundance above 3% are displayed.

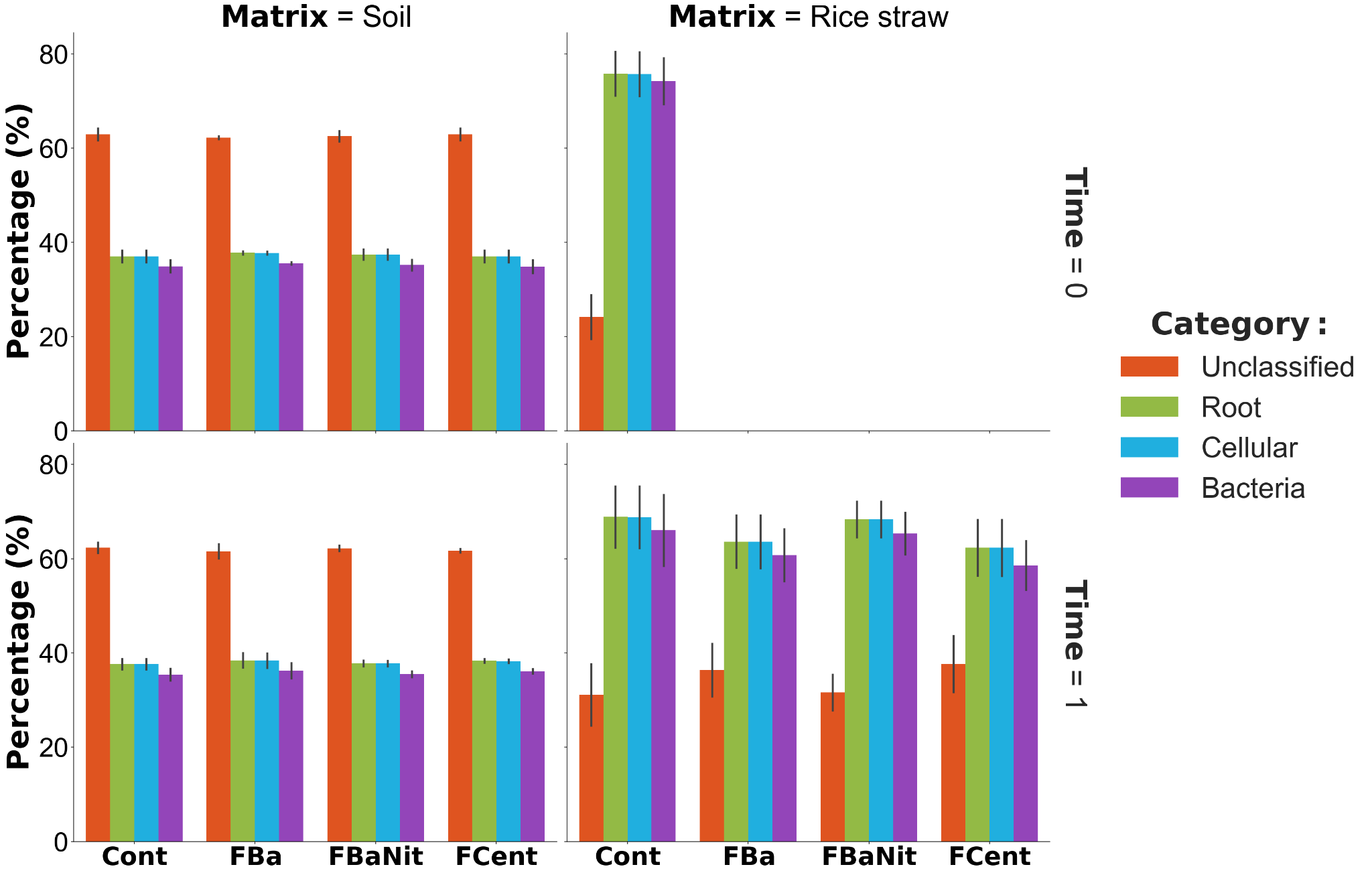

***Fig. S9.*** Percentage of annotated reads using Kraken2/Bracken segmented by matrix and time point. For RS at time *0* samples were taken from a bulk of material; this same bulk was subsampled with specific amounts of RS to start the experiments, and hence accounting with only 1 sample (5 replicates) at time *0*.

**MAG reconstruction and annotation**

A first glimpse into the presence/absence of the different MAG taxonomic annotation across samples allows to capture the general patterns and presumed effects of the treatments. Particularly, in RS samples (***Fig. S13***), the MAGs of an important genus known for its cellulolytic capacity, *Cellulomonas*, were reported only in the *Cont* samples at time *0* and in *FBaNit* samples at time *1*. Also, some representative genera seemed to be affected by the elapsed reaction of RS degradation given that they disappear when comparing against *Cont* samples at time *0*, or at least their abundance was decreased enough for not being assembled, such as *Lactococcus*, *Pantoea*, *Belnapia*, *Bosea*, *Thermomonas*, among others; *Acinetobacter* catches special attention since its MAG abundance was above 20% at time *0*, and after treatment application, it was not observed in treatments *FBa* and *FBaNit*, while in treatment *FCent* its abundance was drastically diminished. On the other hand, *Terrococcus* sequences were abundant enough to build MAGs in all treatments at time *1*, considering this genus was not recovered at time *0*.

Furthermore, the genera *Rhodococcus*, *Enterococcus*, *Melittangium*, *Pararcticibacter*, *Nannocystis*, *Pseudorhodoferax* and *Sphingobium* exhibited a similar trend given the fact that they were not assembled as MAGs after 30 days of treatments. On the contrary, in all treatments (*FBa*, *FBaNit*, *FCent*) the rise of *Nakamurella* and *Agrococcus* seemed to have been encouraged. Important to mention the case of *Pantoea* since the MAGs classified in this genus were abundant in more than 20% at time *0*, although in no case these same MAGs were retrieved at time *1*. However, the effect of the treatments, not limited to, can be observed in the following situations: *i)* the treatment *FBa* seemed to promote the growth of genera such as *Actinomycetospora*, *Actinosynnema*, *Algoriphagus*, *Dyadobacter*, *Hyphomicrobium*, *Longimicrobium*, *Pseudorhodoplanes*, and *Rhodopseudomonas*; *ii)* the nitrogen applied in the treatment *FBaNit* appeared to enable the recovery of *Fimbriimonas*, *Amaricoccus*, *Avipropionibacterium*, *Micromonospora*, *Nesterenkonia*, *Patulibacter*, *Pseudokineococcus*, *Spirosoma*, and *Nocardiopsis*, this former genus represented around 20% of the MAG abundance for this treatment; and *iii)* in treatment *FCent* it seems that the growth of *Methylobacterium* was prevented, in turn that it appeared to stimulate the presence of *Litorihabitans*, *Chryseobacterium*, *Aliihoeflea*, *Aquincola*, and *Agrobacterium*, an important genus in the field of plant molecular biology given its ability related to horizontal transfer genes. Further, solely *Mycobacterium* and *Ochrobactrum* were recovered as MAGs in all treatments, and at time *0* and time *1*, hence representing the core abundant MAGs.

In the case of MAGs from the soil matrices (***Fig. S14***), the annotation provided at genus level is not fully informative since many of the genera names are cryptographic sequences representing genomes that are not fully characterized within the GTDB. Beyond this, given the soil complexity previously mentioned, unclassified MAGs at genus level represent more than the 30% of the MAGs recovered in all samples. Some peculiar cases are illustrated by the appearance of the genera *JAJVID01* (p__Acidobacteriota), *SXND01* (p__Tectomicrobia), *SZUA-320* (p__Gemmatimonadota)*, VAZQ01* (p__Pseudomonadota) and *UBA5189* (p__Chloroflexota) only in *FBaNit* treatment at time *1*. In addition, the genera *JACCXG01* and *Gp7-AA6*, part of the Acidobacteriota phylum, were built only in *Cont* and *FCent* samples, respectively, after 1 month. Given that these genomes are still in study, their function and role within the community are yet to be determined. On the other hand, in terms of the core abundant MAGs, the soil ecosystem appeared to have a strong prevalence of the phylum *Actinomycetota* given that at least 4 of its genera were identified in all treatments at the beginning and the end of the experiments. This core abundant MAGs is completed by *Nitrososphaera*, a genus derived from the domain *Archaea*, that encompasses species able to oxidize ammonia.

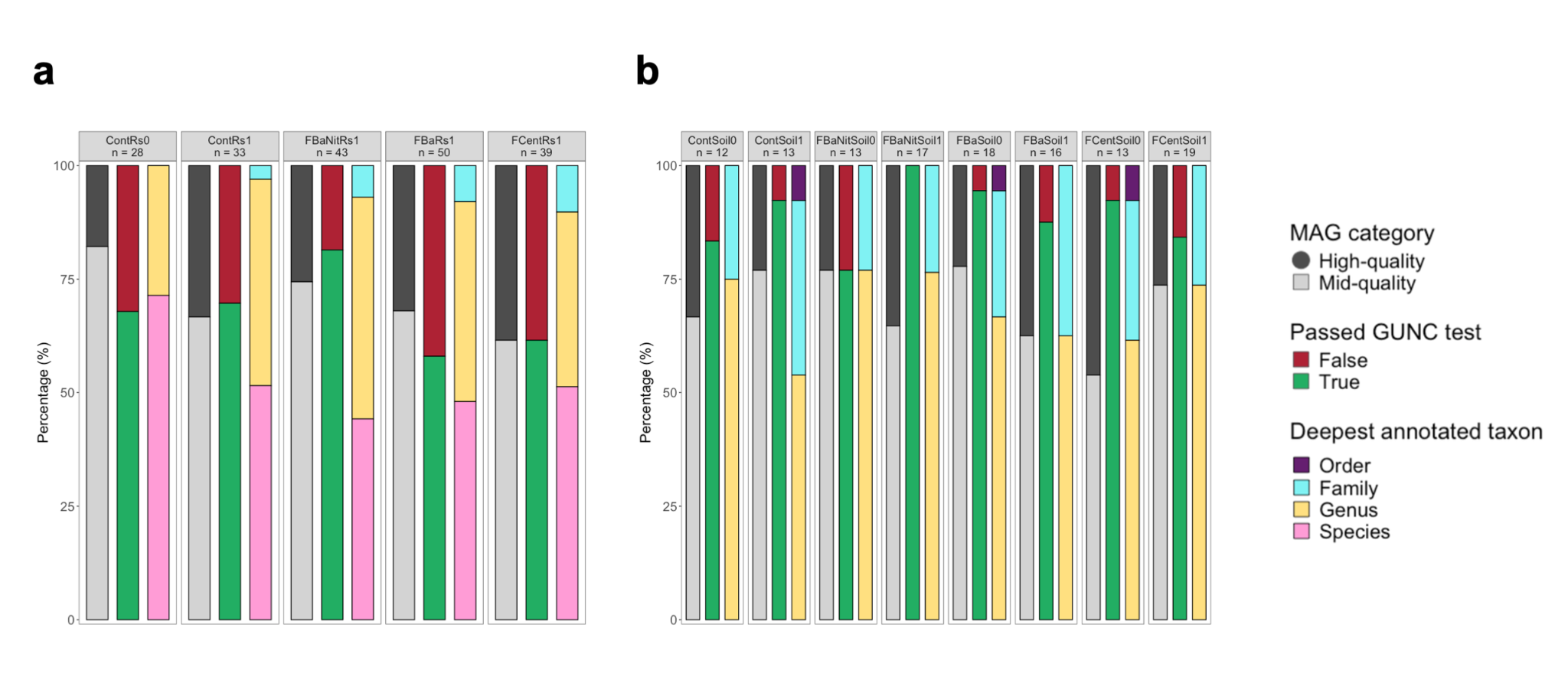

***Fig. S10***. Metrics summaries of the MAGs built from **RS** (***a***) and **soil** (***b***) samples; the columns represent, in order, the proportion of: mid-quality and high-quality MAGs according to CheckM2 estimations; MAGs passing the GUNC test; and annotated MAGs at different taxonomic levels based on GTDB-Tk2.

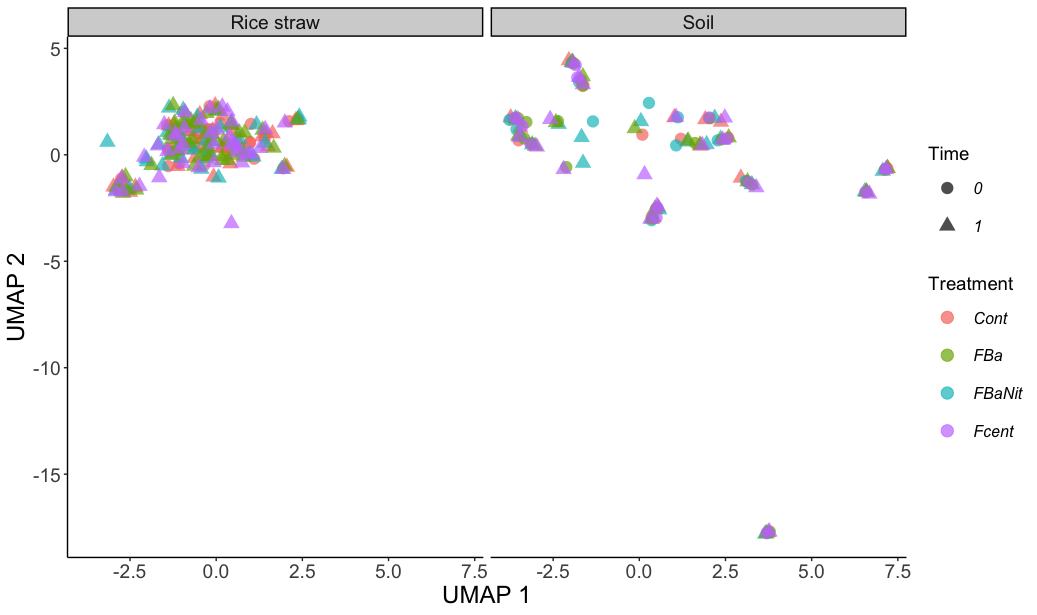

***Fig. S11***. Two-dimensional Uniform Manifold Approximation and Projection (UMAP) embedding of normalized CAZy enzymes counts per MAG (before clustering with FastANI) colored by treatment and shape by the time in which the MAGs were recovered.

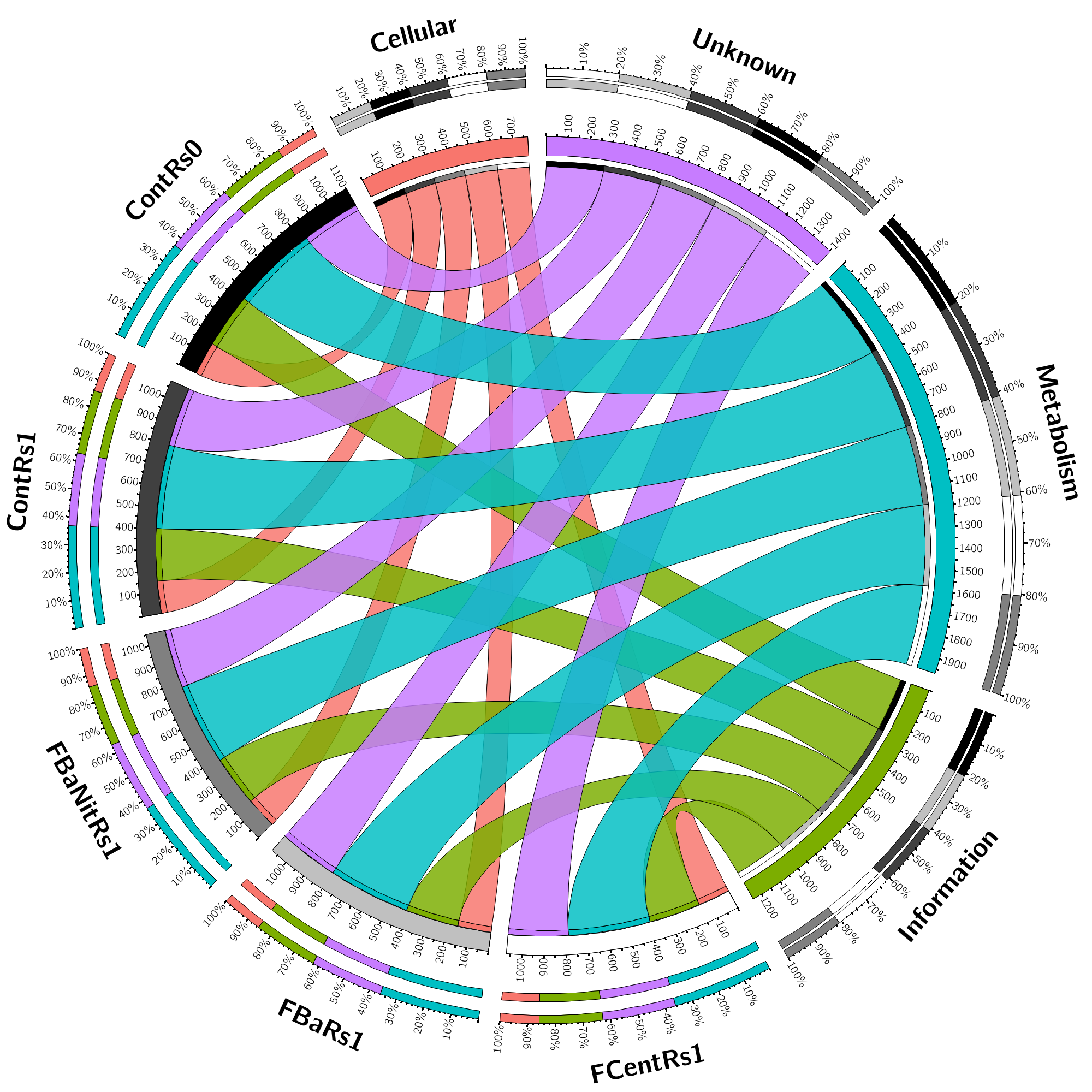

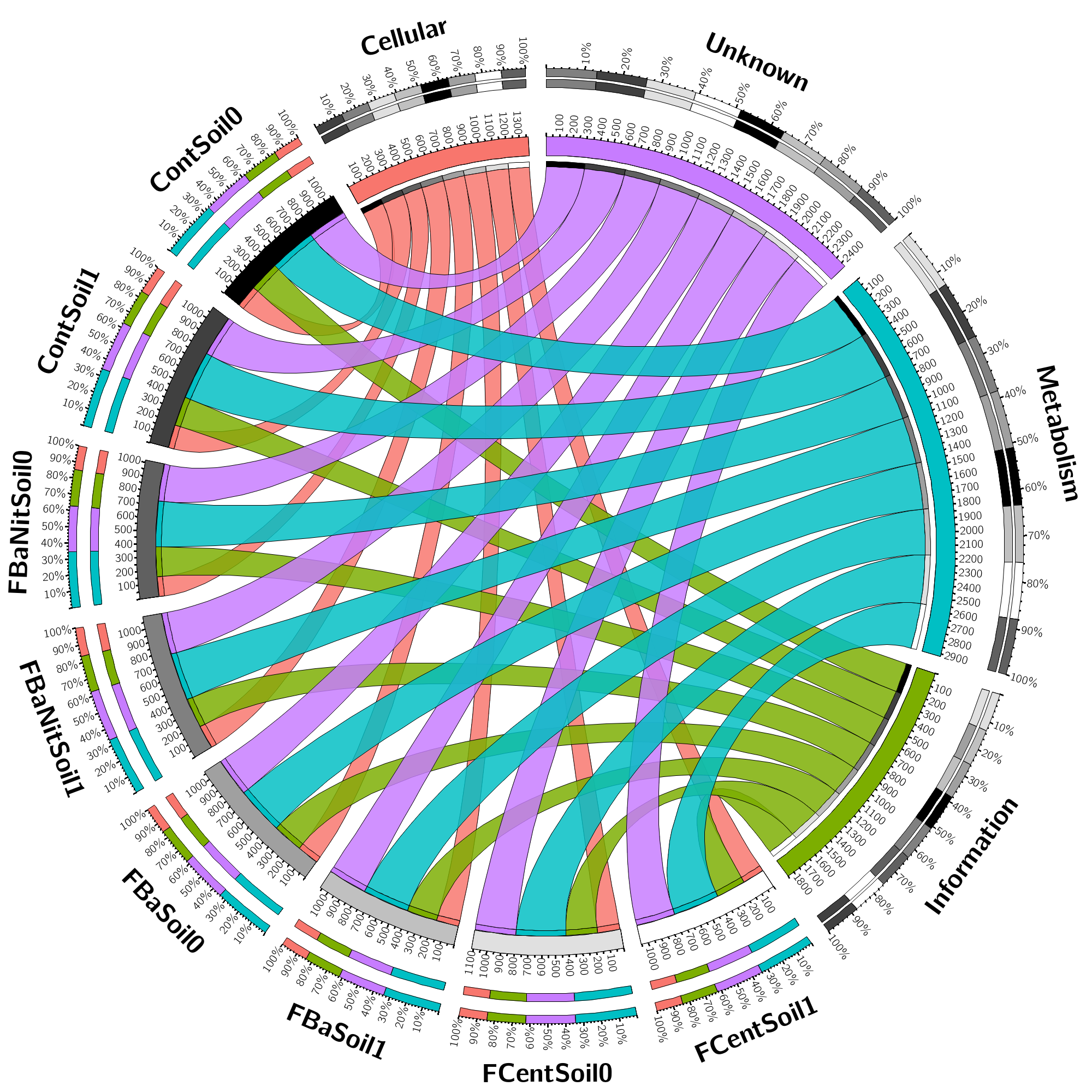

***Fig. S12***. Circos plots showing the proportion of the 4 main categories (Metabolism; Information Storage and Processing; Cellular Processes and Signaling; and Protein Coding Sequences with Unknown function) of Clusters of Orthologous Genes (COGs) found within the MAGs recovered from **RS** (***top***) and **soil** (***bottom***) the samples. The inner ring accounts for the total number of COGs related to every sample or COG category; the outer ring depicts the relative abundance of COGs from each sample or COG category; the width of the ribbons linking any sample and a COG category depicts their relative abundance to each other. The number of counts per category was normalized using the total number of genes found per MAG.

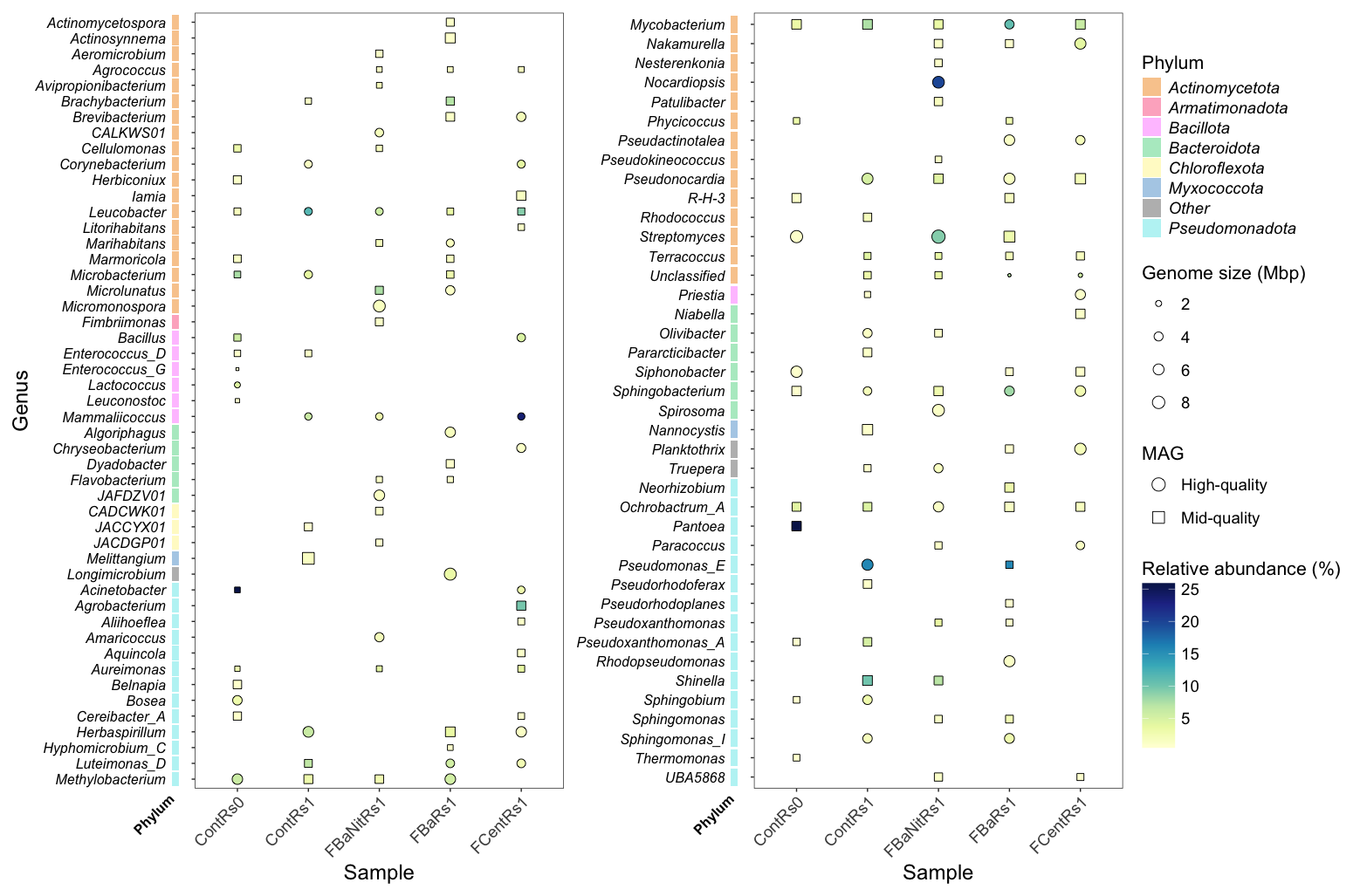

***Fig. S13***. Presence/absence of MAGs annotated with GTDB-Tk2 in **RS** samples at genus level. Genome size, in Mbp, is indicated as the point size, while the color scale represents the relative abundance of each MAG per sample, and the point shape shows whether the displayed MAG is classified as high-quality or mid-quality (CheckM2). When MAGs with the same genus annotation were found in the same sample, the most complete one is depicted and their abundances are added up.

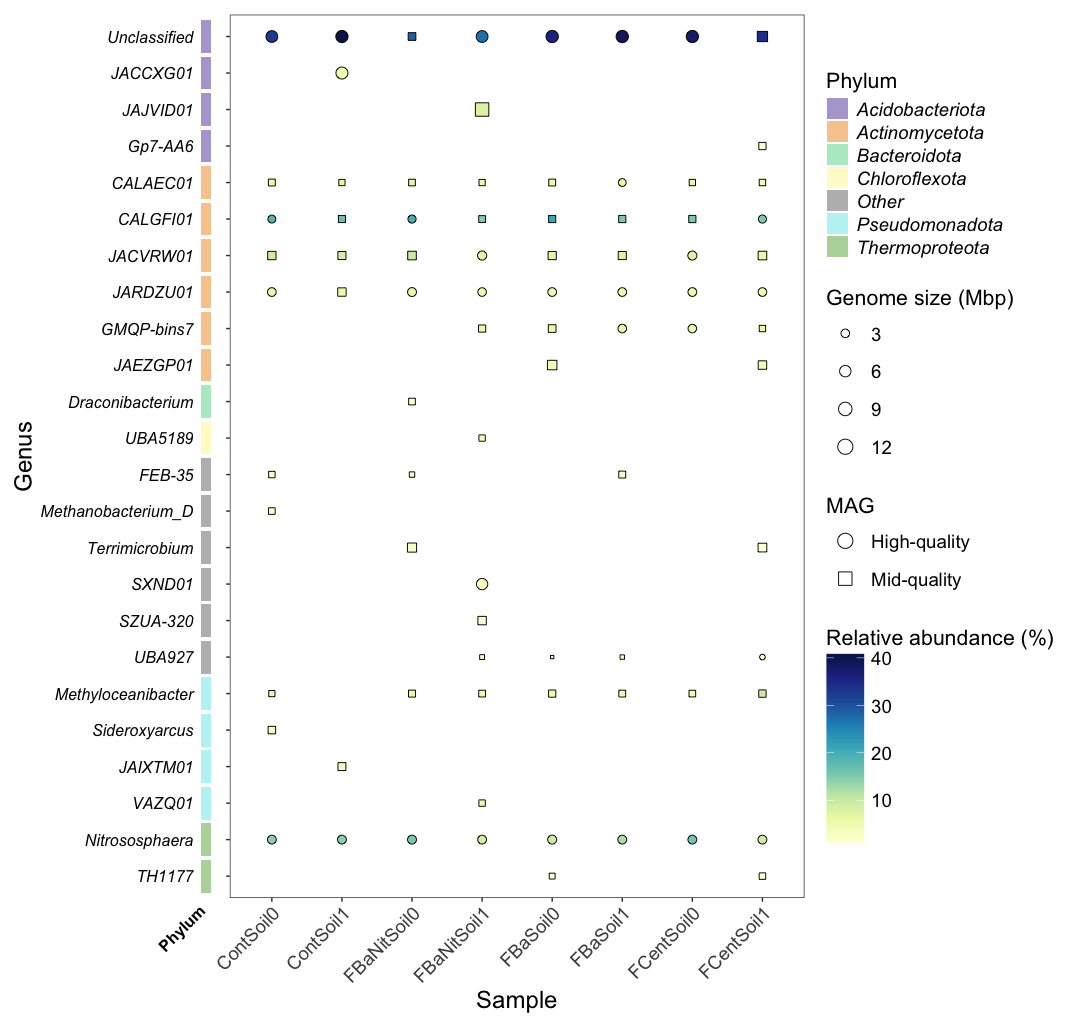

***Fig. S14***. Presence/absence of MAGs annotated with GTDB-Tk2 in **soil** samples at genus level. Genome size, in Mbp, is indicated as the point size, while the color scale represents the relative abundance of each MAG per sample, and the point shape shows whether the displayed MAG is classified as high-quality or mid-quality (CheckM2). When MAGs with the same genus annotation were found in the same sample, the most complete one is depicted and their abundances are added up.

**MAGs and integration of carbohydrate-active enzyme information**

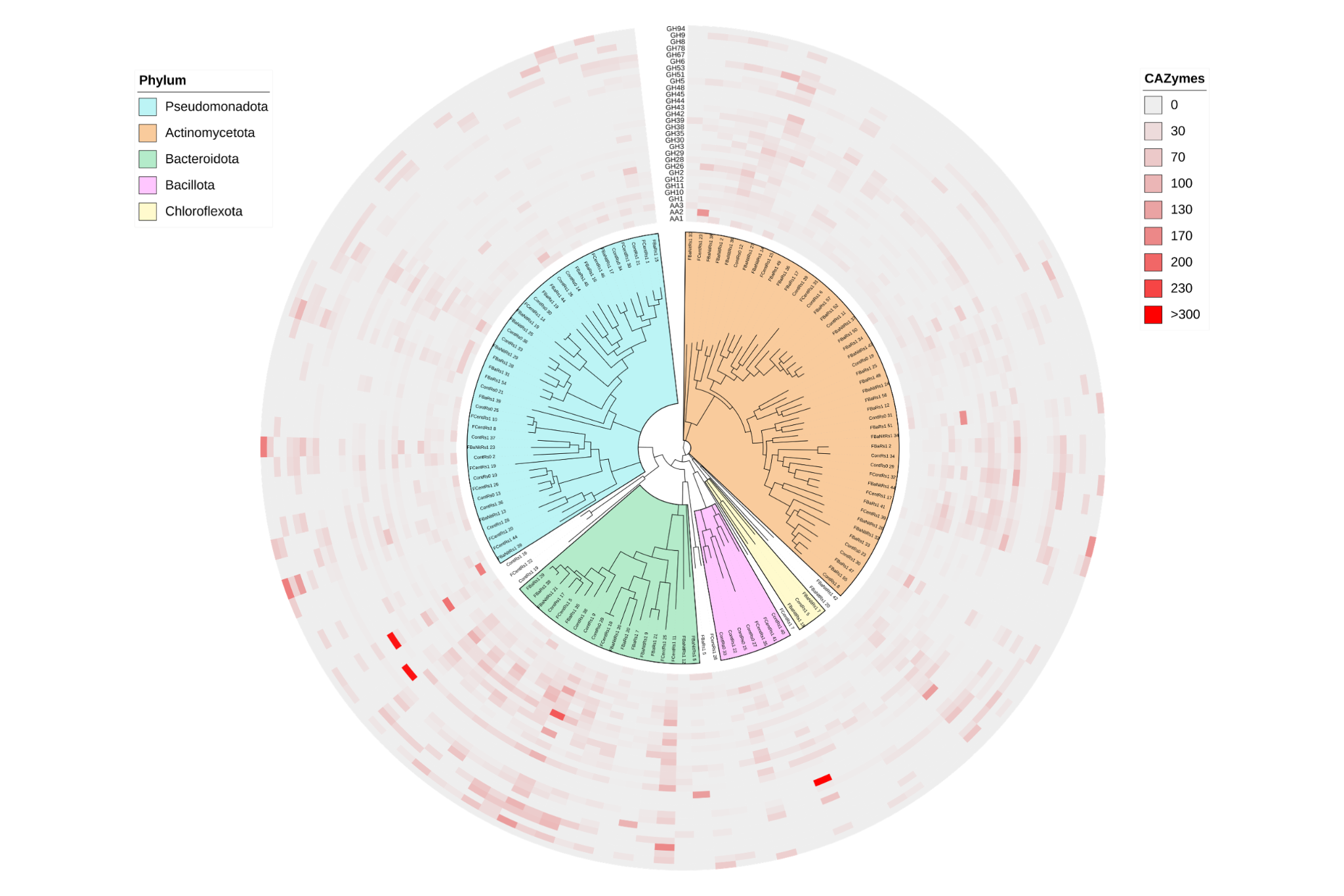

***Fig. S15***. 125 clustered MAGs recovered from **RS** samples and their normalized counts for CAZy families associated with lignocellulose-degrading enzymes. The organization of the tree relies on the phylogenetic information retrieved from GTDB-Tk2. The displayed MAGs are representatives from the generated clusters using ANI, and they were selected based on the following score: completeness – 0.5 x contamination.

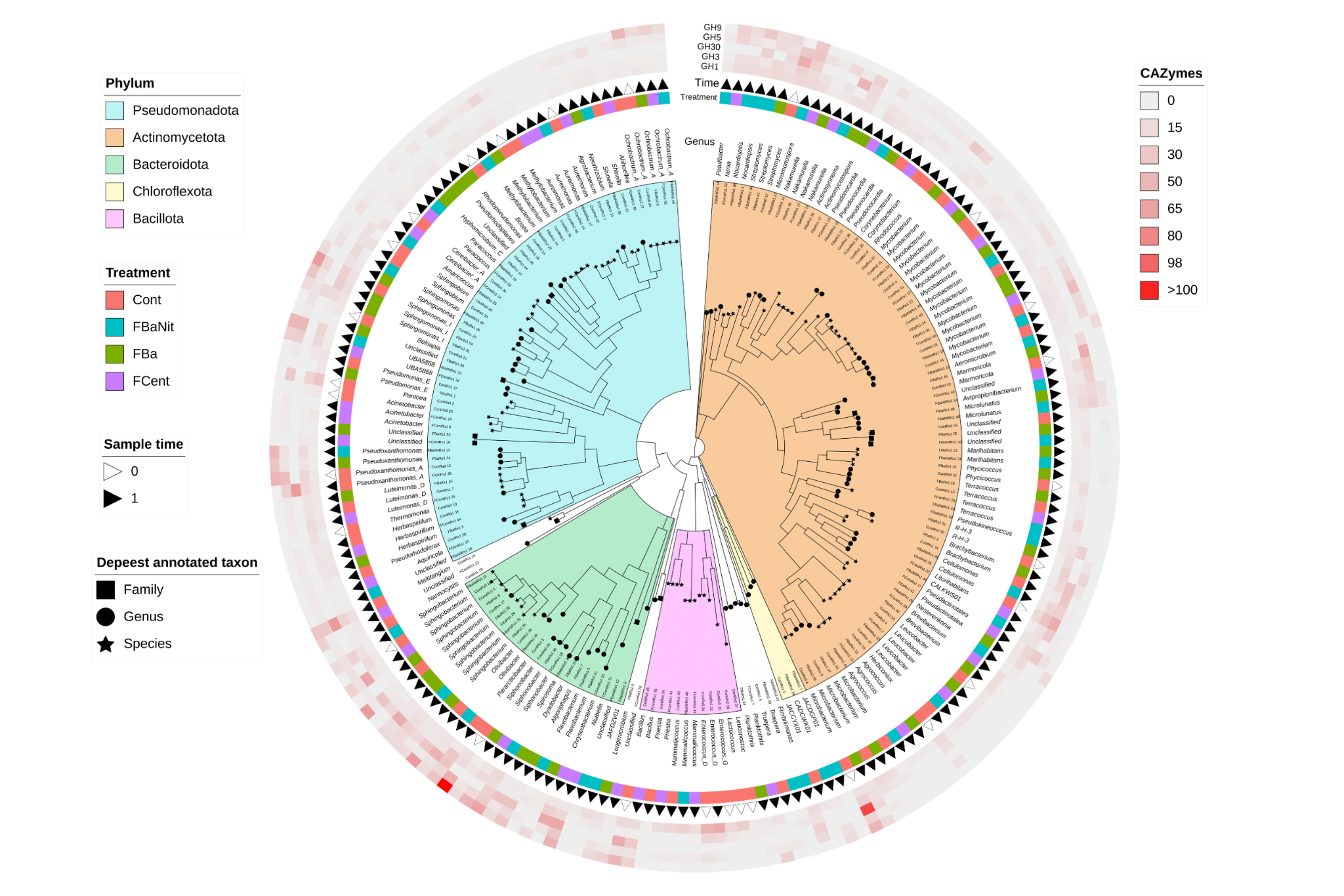

***Fig. S16***. 193 unclustered MAGs recovered from **RS** samples and their normalized counts for CAZy families associated with β-glucosidases. The organization of the tree relies on the phylogenetic information retrieved from GTDB-Tk2. The color strip indicates the treatment matrix in which each MAG was built, and the triangle represents at which time the MAG was found. The visualization also includes genus annotation, Phylum affiliation and the deepest annotated taxon per MAG.

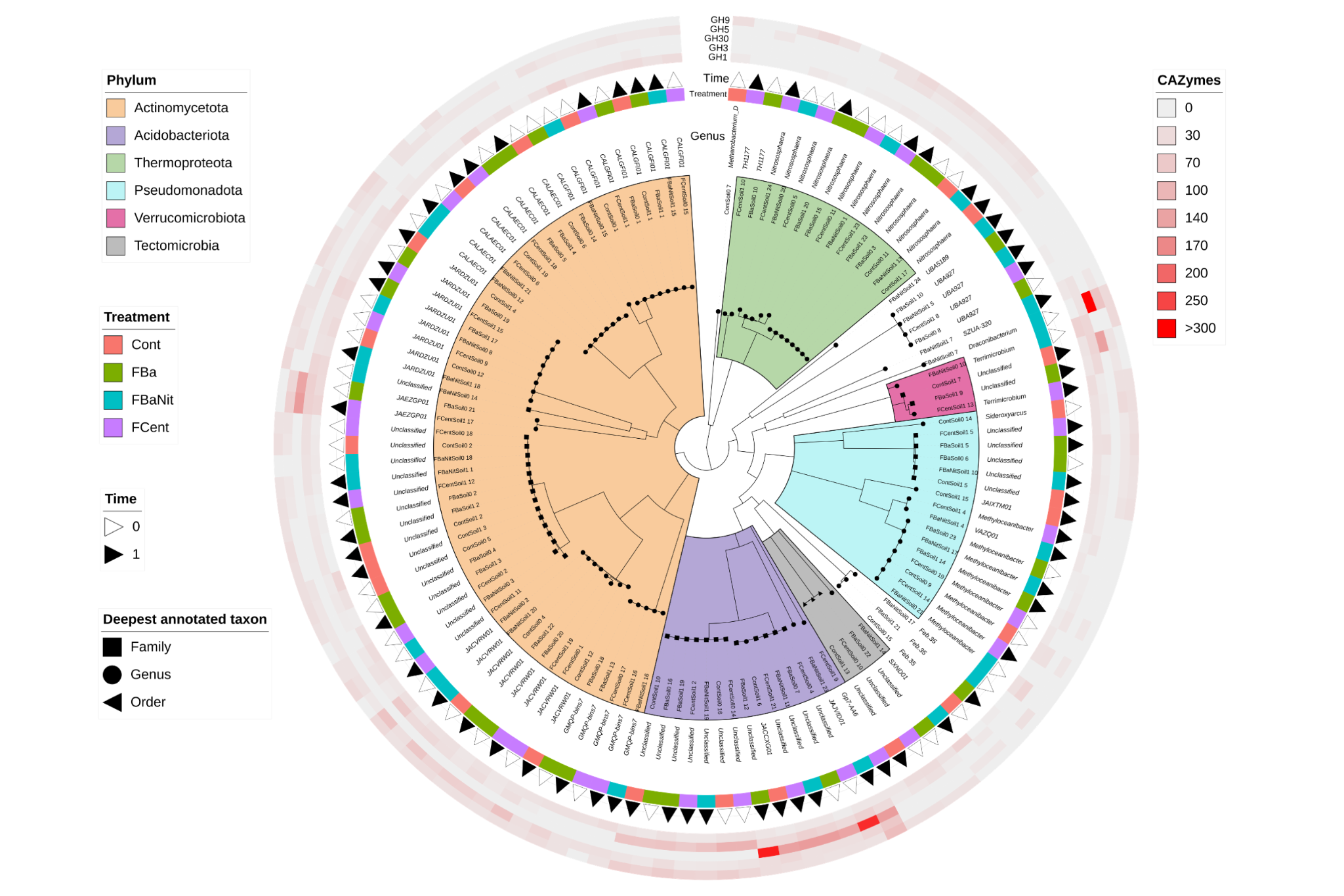

***Fig. S17***. 121 unclustered MAGs recovered from **soil** samples and their normalized counts for CAZy families associated with β-glucosidases. See ***Fig. S16*** for feature explanation.

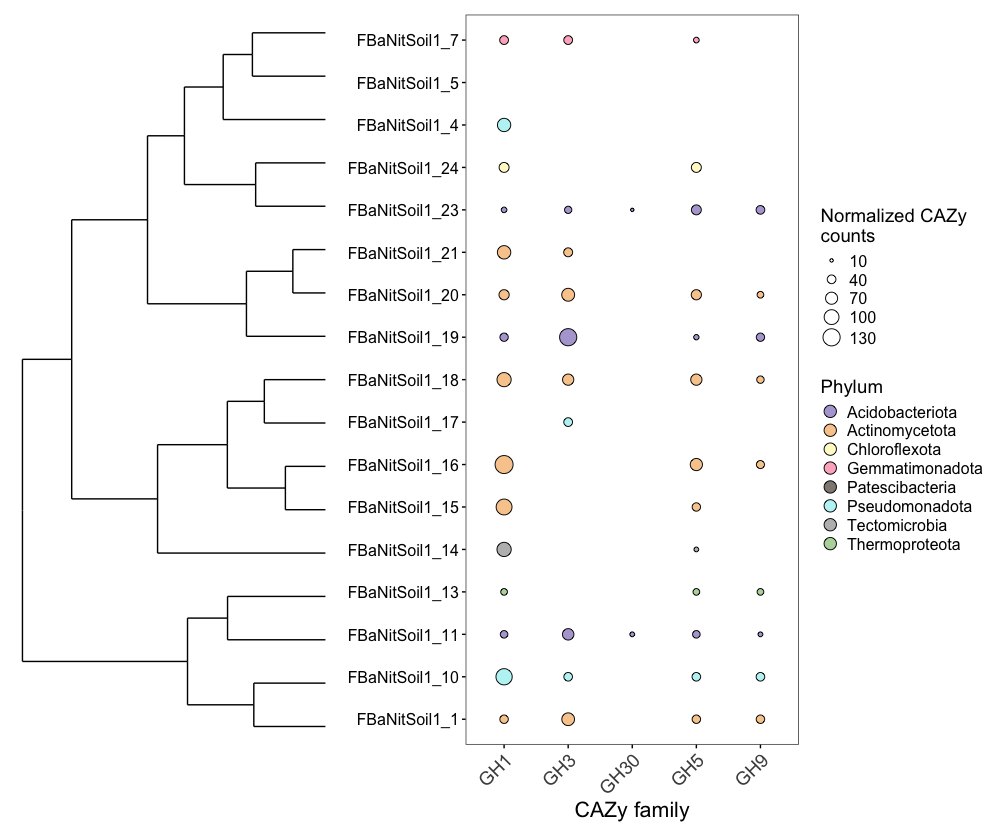

***Fig. S18***. Bubble plot depicting the number of normalized counts for CAZyme families associated with β-glucosidases found in the MAGs recovered from samples ***FBaNit1*** in soil. Visualization includes: MAG clustering based on normalized CAZyme counts of the selected families, and phylum information (GTDB-Tk2).

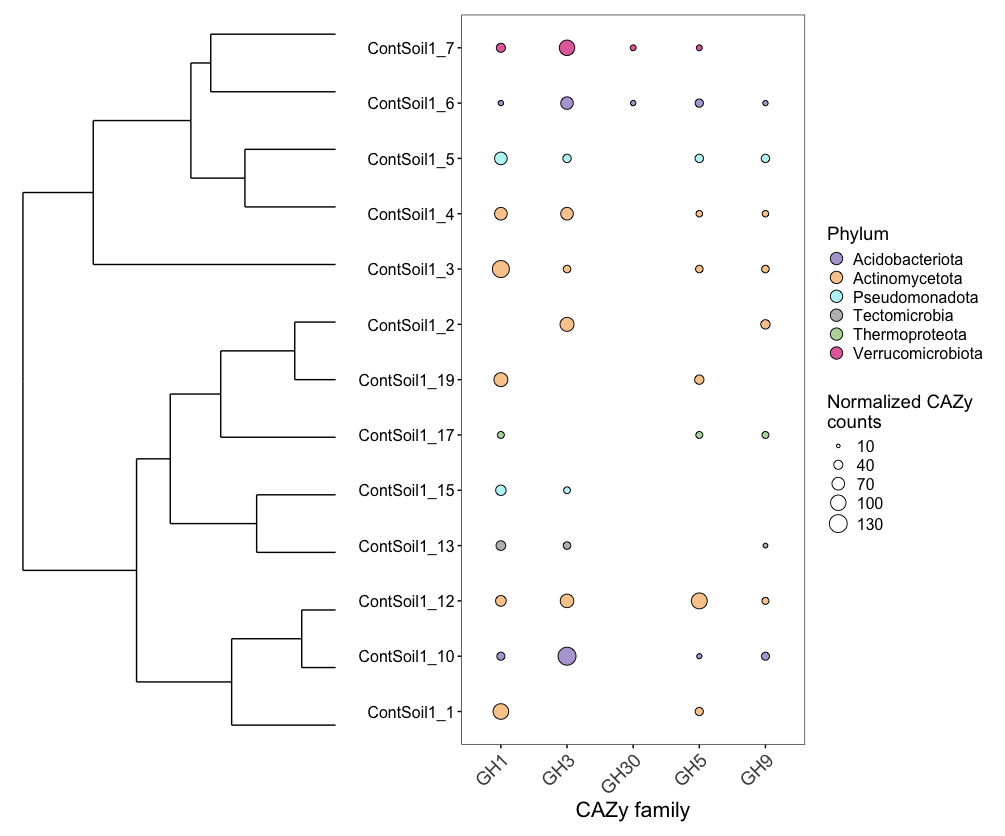

***Fig. S19***. Bubble plot depicting the number of normalized counts for CAZyme families associated with β-glucosidases found in the MAGs recovered from samples ***Cont1*** in soil. Visualization includes: MAG clustering based on normalized CAZyme counts of the selected families, and phylum information (GTDB-Tk2).
